# Alternative futures for global biological invasions

**DOI:** 10.1101/2021.01.15.426694

**Authors:** Núria Roura-Pascual, Brian Leung, Wolfgang Rabitsch, Lucas Rutting, Joost Vervoort, Sven Bacher, Stefan Dullinger, Karl-Heinz Erb, Jonathan M. Jeschke, Stelios Katsanevakis, Ingolf Kühn, Bernd Lenzner, Andrew M. Liebhold, Michael Obersteiner, Anibal Pauchard, Garry D. Peterson, Helen E. Roy, Hanno Seebens, Marten Winter, Mark A. Burgman, Piero Genovesi, Philip E. Hulme, Reuben P. Keller, Guillaume Latombe, Melodie A. McGeoch, Gregory M. Ruiz, Riccardo Scalera, Michael R. Springborn, Betsy von Holle, Franz Essl

## Abstract

Scenario analysis has emerged as a key tool to analyze complex and uncertain future socio-ecological developments. However, current global scenarios (narratives of how the world may develop) have neglected biological invasions, a major threat to biodiversity and the economy. We used a novel participatory process to develop a diverse set of global biological invasion scenarios spanning a wide breadth of plausible global futures through 2050. We adapted the widely used “two axes” scenario analysis approach to develop four families of four scenarios each, resulting in 16 scenarios that were later clustered into four contrasting sets of futures. Our analysis highlighted that socio-economic developments and changes in sustainability policies and lifestyle have the potential to shape biological invasions, in addition to well-known ecological drivers, such as climate and human land use change. Our scenarios align fairly well with the recently developed shared socio-economic pathways, but the factors that drive differences in biological invasions are underrepresented there. Including these factors in global scenarios and models is essential to adequately consider biological invasions in global environmental assessments, and obtain a more integrative picture of future socio-ecological developments.

## Introduction

Biological invasions are recognized as a major driver of biodiversity loss (Pyšek et al. 2020). Invasive alien species (IAS) can threaten native biota and alter ecosystem functioning, disrupt the delivery of ecosystem services and cause numerous social and economic impacts (Diagne et al. 2020; Simberloff et al. 2013). The number of alien species continues to increase globally with no sign of saturation despite efforts to halt invasions (Seebens et al. 2017). Advances have been made in understanding the direct drivers of biological invasions, such as ecological determinants (Simberloff et al. 2013) and anthropogenic factors including climate change (Hulme 2017), global trade (Early et al. 2015) and human disturbance (Spear et al. 2013). However, it is still unclear how these direct drivers are shaped by social developments and how indirect social drivers may determine the future influence of biological invasions (e.g. Lotz and Allen 2013). A better understanding of how social change may determine biological changes is a prerequisite to understand and effectively manage, biological invasions in the Anthropocene (Essl et al. 2020).

Scenario analysis provides a systematic method to assess how complex interactions among many drivers of change may produce multiple plausible futures (Peterson et al. 2003). Scenario analysis has been increasingly used to analyze likely outcomes of global and regional environmental developments (Oteros-Rozas et al. 2015; Pereira et al. 2020; Spangenberg et al. 2012; van Vuuren et al. 2014). Scenarios are neither predictions nor forecasts, rather they are descriptions and/or qualitative explorations of alternative paths along which the future might unfold (Van der Heijden 2005). The comparative analysis of a set of scenarios can be used to identify key uncertainties, and allows the incorporation of alternative or competing perspectives and theories into analysis of potential futures (Peterson et al. 2003). Scenarios are qualitative in nature, although they can be combined with models to produce quantitative estimates of future changes (Lenzner et al. 2019).

We used scenario analysis to investigate the complex and uncertain interactions underlying biological invasions, and to capture a variety of expert knowledge on how biological invasions interact with other relevant drivers of global change. The development of scenarios allowed us to explore a wide range of potential variations in the number of IAS that are likely to become established through 2050. While scenario-based analyses of biological invasions have been undertaken at regional scales (e.g. Chytrý et al. 2012; Le Maitre et al. 2004; Roura-Pascual et al. 2011), the last global scenario analyses incorporating biological invasions were done over two decades ago (Carpenter et al. 2005; Sala et al. 2000), and focused on drivers of biodiversity change rather than invasions themselves. Therefore, undertaking a science-based analysis of global trajectories of biological invasions both addresses a critical research gap (IPBES 2016), and is a timely contribution to the future quantification of the effects of biological invasions on the environment and human livelihoods (Lenzner et al. 2019). We developed new global scenarios for biological invasions to avoid being constrained by pre-existing scenarios that were not created with a focus on the drivers of IAS change, such as the widely used Shared Socio-Economic Pathways (SSPs). The SSPs global change scenarios were developed by the climate change research community and include both qualitative descriptions and quantifications of broad trends in socio-economic developments (van Vuuren et al. 2014). They serve as a basis for integrated assessment models simulating the evolution of land use, energy consumption and resulting greenhouse gas emissions under different SSPs and climate policy assumptions (O’Neill et al. 2017). However, the development of these scenarios has primarily focused on climate change and they only offer one set of scenarios, framed by a specific set of assumptions, excluding many other framings of potential futures. We nonetheless compared the SSPs against our biological invasion scenarios, to advance the integration of invasion science into global environmental assessments, a topic that has been identified as a research priority (CBD 2010; IPBES 2016)..

## Methods

### Scenarios development

We adapted the common “two axes” scenario analysis approach (Van der Heijden 2005) to develop four families of four scenarios each. While the two-axes scenario approach has strong benefits in its ability to communicate a set of scenarios quickly and in a transparent manner, the use of a single set of scenarios framed through this approach has been criticized, because it limits the exploration of futures through a single set of assumptions and drivers (Vervoort et al. 2015). There are alternative methods that integrate many drivers in a single scenario set, but such approaches end up with a limited set of plausible futures, and run the risk of being more opaque in their assumptions (Lord et al. 2016).

Our approach was to construct multiple two-axis scenario sets – this approach allowed us to investigate potential futures through multiple framings based on different driver combinations. This, in turn, enabled us to explore a much more multidimensional possibility space for plausible trajectories of biological invasions. We then clustered the 16 scenarios we had developed into a reduced and manageable set of futures (representing archetypes of scenarios) to facilitate comparisons with the single set of widely used SSPs (Fig. 1).

**Fig. 1.**
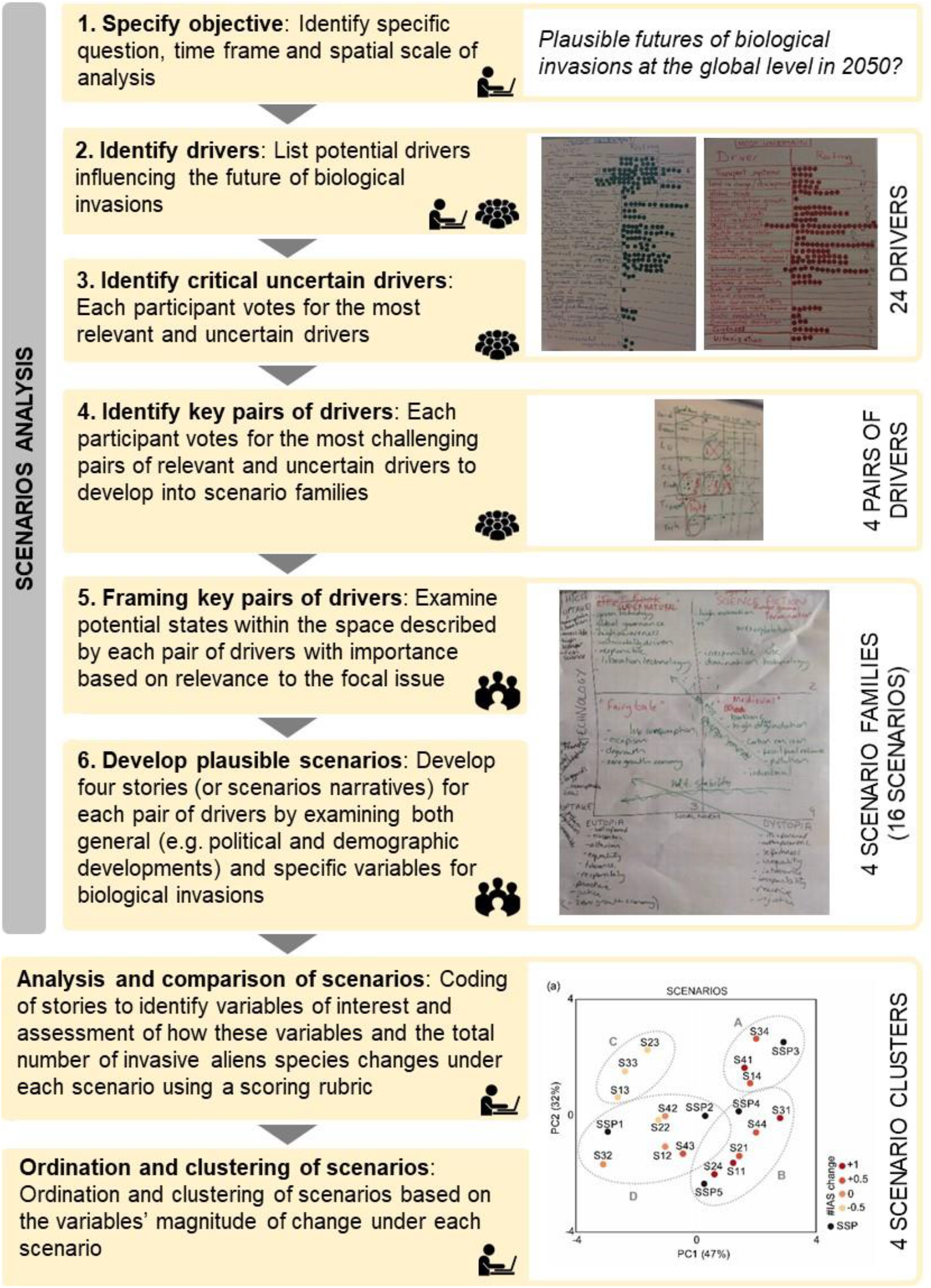
Flow chart and photographs showing methodologies used to develop global scenarios for biological invasions through 2050. The icons indicate the type of activity and the people involved in each step (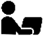: online contributions of all participants, 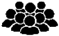: contributions of workshop participants, 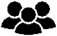: contributions of breakout groups).

The scenario analysis was conducted using a six-step participatory process, combining expert-based online discussions with a two-day workshop (Vienna, 6-7 October 2016) (Fig. 1). The participants were mainly invasion biologists (24 out of 30), but there were also four experts on topically relevant research such as global change biology and environmental economy and two others on scenario approaches (Table S1). Attendants to the workshop (19) were mainly from Europe due to the nature of the funding body (COST Action TD1209: ALIEN Challenge), but there were also three non-European experts. The remaining of participants that contributed online before and after the workshop were four European participants and seven international collaborators (Table S1). All participants are authors of the manuscript.

#### Step 1. Specify objective

All participants (30) agreed upon the objective of the scenario analysis through online discussions, which was: “Exploring different plausible futures concerning biological invasions at the global level through 2050”.

#### Step 2. Identify drivers

Identification of drivers influencing biological invasions through 2050. Scenario experts (L. Rutting and J. Vervoort) reviewed a wide range of existing scenario exercises (Alexandratos and Bruinsma 2012; CCAFS 2014; Gallopin et al. 1997; IPCC 2000, 2013; Mora et al. 2016; OECD 2009; Palazzo et al. 2016; Rockefeller Foundation and GBN 2010; Vervoort et al. 2016; Vervoort et al. 2013) and identified the drivers considered in these future scenarios. This preliminary list of drivers was updated (by adding/removing/modifying certain drivers) by all participants prior to the workshop, but also during the scenario workshop taking into account likely influences of these drivers on biological invasions.

#### Step 3. Identify critical uncertain drivers

Participants at the workshop selected the most relevant (i.e. important for biological invasions) and uncertain drivers (i.e. how unknown is the range of plausible different directions this driver may take) from the list drawn in the previous step based on a voting system. Each participant had 10 points to use to rank the most relevant drivers influencing the future status of biological invasions, and an additional 10 points to rank the most uncertain drivers based on their expertise. Scores were distributed freely among the drivers, with the possibility to assign multiple points to a single driver. The drivers were then plotted in a coordinate system on the basis of their relevance and uncertainty. Workshop participants examined the position of each driver in the coordinate system and selected through group discussion the highest ranking drivers.

#### Step 4. Identify key pairs of drivers

This step involved the selection of pairs of drivers that led to the most challenging, diverse and relevant scenarios for exploring the future of biological invasions. Instead of choosing one single pair of drivers (Van der Heijden 2005) or combining a larger number of drivers (Lord et al. 2016) into one overarching, small set of scenarios, workshop participants discussed and voted on the most useful (i.e. challenging, diverse and relevant) pairs of two drivers. First participants voted for all possible pairs of drivers retained in the previous step and identified the pairs with the most votes, and then voted and selected the four most useful pairs among those with the most votes obtained in the previous voting round. In each voting round, each participant at the workshop had 5 points to distribute freely among the pairs of drivers; it was possible to assign multiple points to a single pair or to leave some points unassigned. These four pairs of drivers were developed into four scenario families (composed of four scenarios each), and each pair of drivers determined the framing of each future in the family in the following steps.

#### Step 5. Framing key pairs of drivers

Examination and framing of each of these four pairs of drivers was carried out by breakout groups composed of 4-5 workshop participants, who discussed the different possible states of the driver axes. Drivers can be conceptualized in different ways, for instance, economic development can be defined along extremes of high or low, stable or unstable, equal or unequal, etc. The participants in the workshop defined which set of driver states was most useful for exploring futures of biological invasions. They did so by examining combinations of driver states and what types of scenarios these combinations yield – if a combination did not yield useful (challenging, diverse, relevant) scenarios, then they considered a different combination.

#### Step 6. Develop plausible scenarios

Scenarios for biological invasions were developed by the same breakout groups that in the previous step examined and framed for each pair of drivers. Each pair of drivers formed a pair of axes, which yielded four scenarios. The group developed narratives for these scenarios (i.e. descriptions of how the future may unfold under each scenarios), by examining both general, contextual developments (i.e. political and demographic developments; socio-economic and trade developments; lifestyle and values; technological developments; and environmental developments and natural resources) and specific details on biological invasions. It is important to note that while selected pairs of drivers served as starting points and a way to frame the scenarios, the other drivers from step 2, which did not define the scenario axes, could also be taken into consideration in the development of the scenario narratives themselves. Since each group worked on one pair of drivers, the workshop resulted in four families of four scenarios for a total of 16 different scenarios.

### Scenarios comparison

To identify similarities and differences among the scenarios for biological invasion created during the scenario analysis, but also between these scenarios and the widely used SSPs, we clustered all scenarios based on a set of variables. To select these variables, first we coded the scenarios narratives by means of qualitative content analysis. We identified terms that symbolically captured the essence of the different portions of the narratives and organized them in categories (Saldaña 2013). These terms (hereafter referred to as variables) are affected by variations in scenario assumptions, but do not necessarily correspond to the drivers used to frame the scenarios. We identified those variables that appeared in three or four families of scenarios and selected them as the variables of interest to compare the scenarios.

Then, we qualitatively assessed how these variables of interest were likely to change under different scenarios for biological invasions by 2050, as well as the resulting total number of IAS. The assessment of these variables and the number of IAS was standardized using a scoring rubric, where the magnitude of change was measured using a 5-level Likert scale ranging from +1 (high increase) to −1 (high decrease) (Table S3). A value of 0 (no change) designates the current rate of change, while “increase” implies an acceleration and “decrease” implies a slowing of the current rate of change. This scoring rubric was created by the study participants with the specific purpose to compare the scenarios. To facilitate the assessment and to ensure equivalencies among the different ratings, each level had a description associated to it. We attempted to link these descriptions to existing publications assessing and/or projecting changes in future trends related to each variable (i.e. publications embedded in Table S3); the use of absolute values did not intend to be exact, but served to characterize the magnitude of change of each variable and impacts across each 5designates the current rate level category (Table S3). Workshop participants were asked to assess only the scenarios that they contributed to create, by rating the variables considered in the scoring rubric based on the descriptions of the scenario narratives. This process was conducted online by the four breakout groups that created the scenarios during the workshop (in previous step 6). In addition to the scenarios for biological invasion, we conducted a similar assessment of the SSPs. The assessment of SSPs narratives based on the variables of interest considered in the scoring rubric was based on the scenario descriptions provided in several publications (Calvin et al. 2017; Dellink et al. 2017; Fujimori et al. 2017; Jiang and O’Neill 2017; Kriegler et al. 2017; O’Neill et al. 2017; O’Neill et al. 2016; Riahi et al. 2017; van der Mensbrugghe 2015; van Vuuren et al. 2017).

These expert-based assessments of future changes in variables of interest under the different scenarios were then used to ordinate and cluster the scenarios/variables. First we applied a principal component analysis (PCA) on Spearman covariance matrix using the stats package in R (R_Core_Team 2018), and then used the first two components to visualize the distribution of scenarios in a bivariate plot formed by these two new linear variable combinations. Additionally, we also applied a hierarchical cluster analysis on the scenario and variable coordinates in the main two axes of the PCA (*gplot* package (Warnes et al. 2016) in R). Scenarios for biological invasions were characterized by the number of IAS expected to have become established by 2050 (i.e. last row in Table S3); this variable was not considered in the principal component analysis, but it was used to differentiate scenarios likely to result in a low or high number of IAS, respectively, and to assess the coherence of the scenarios within each cluster. These analyses enabled us to identify the relationships between the scenarios for biological invasions and those of SSP narratives, to ensure the consistency of our scenario narratives but also to contribute to the work of other research programmes assessing the effects of global change on biodiversity.

## Results and Discussion

### Scenarios development

The workshop participants identified 24 drivers as potentially suitable for building the scenarios for biological invasions, which were grouped into five categories: (i) politics and demographics (6 drivers); (ii) economic and trade (3 drivers); (iii) lifestyle and values (4 drivers); (iv) technology (2 drivers); and (v) environment and natural resources (9 drivers) (Fig. 2a). During the workshop, 11 of these 24 drivers were classified as being both relevant and uncertain for the storyline development (Fig. 2b). Although *climate change* (driver 16) was not among the most uncertain drivers, participants decided to include it in the scenario analysis because of its potential to exacerbate invasions (Hulme 2017). Other drivers presenting a high relevance and uncertainty were also combined into a single driver because of their strong relationships. For example, drivers related to the category lifestyle and values (i.e. drivers 11-13) were grouped together, and *economic growth* was merged with *global trade* (drivers 7 and 9, respectively). Seven drivers were finally considered as the most relevant and uncertain ones (Fig. 2b).

**Fig. 2.**
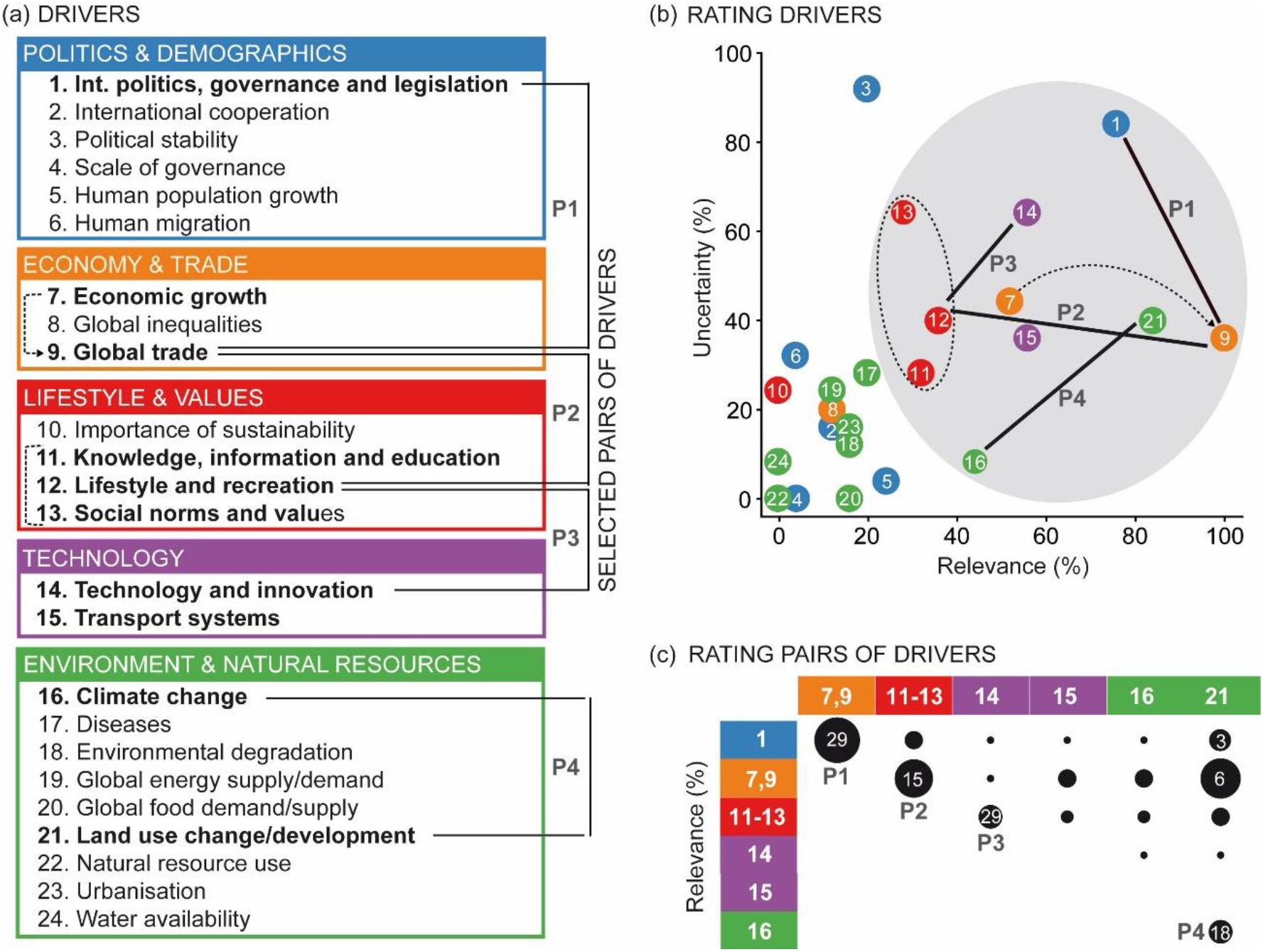
Socio-ecological drivers of biological invasions. (a) list of drivers grouped by categories, (b) rating based on driver’s relevance and uncertainty and (c) rating of selected set of driver pairs. Size of circles in (c) is proportional to the percentage of votes assigned to all driver pairs, while numbers in black circles indicate the percentage of votes given to the final set of driver pairs considered for scenario building. Drivers highlighted in bold in (a) indicate the most relevant and uncertain drivers according to (b). Dashed lines in (a) and (b) indicate drivers that were grouped together, while continuous lines join pairs of drivers used to develop scenarios for biological invasions (abbreviated as P1, P2, P3, P4).

We evaluated all pair-wise combinations of these seven drivers and used voting to select the four pairs of drivers to develop the scenarios. Among all possible pairs of drivers (i.e. 21 pairs), the four pairs considered the most diverging by the participants were: (1) *international politics, governance and legislation* vs. *global trade*; (2) *global trade* vs. *social norms* (composite of drivers 11-13); (3) *social norms* vs. *technology and innovation*; and (4) *climate change* vs. *land use change/development* (Figs. 2c and S1). These pairs of drivers included six unique drivers. These drivers correspond to well-known drivers of biological invasions, such as global trade, land use change/development and climate change, as well as drivers associated to societal variables (usually called indirect drivers, e.g. IPBES (2016)) that have been largely ignored in the invasion literature: international politics, governance and legislation; lifestyle and social norms; and technological development and innovations (Fig. 2c). The latter group of drivers could have an enormous influence on the number of IAS in the future, but they are difficult to quantify. Each of these pairs of drivers was used to create a family of four scenarios (Fig. 3; Text S1).

**Fig. 3.**
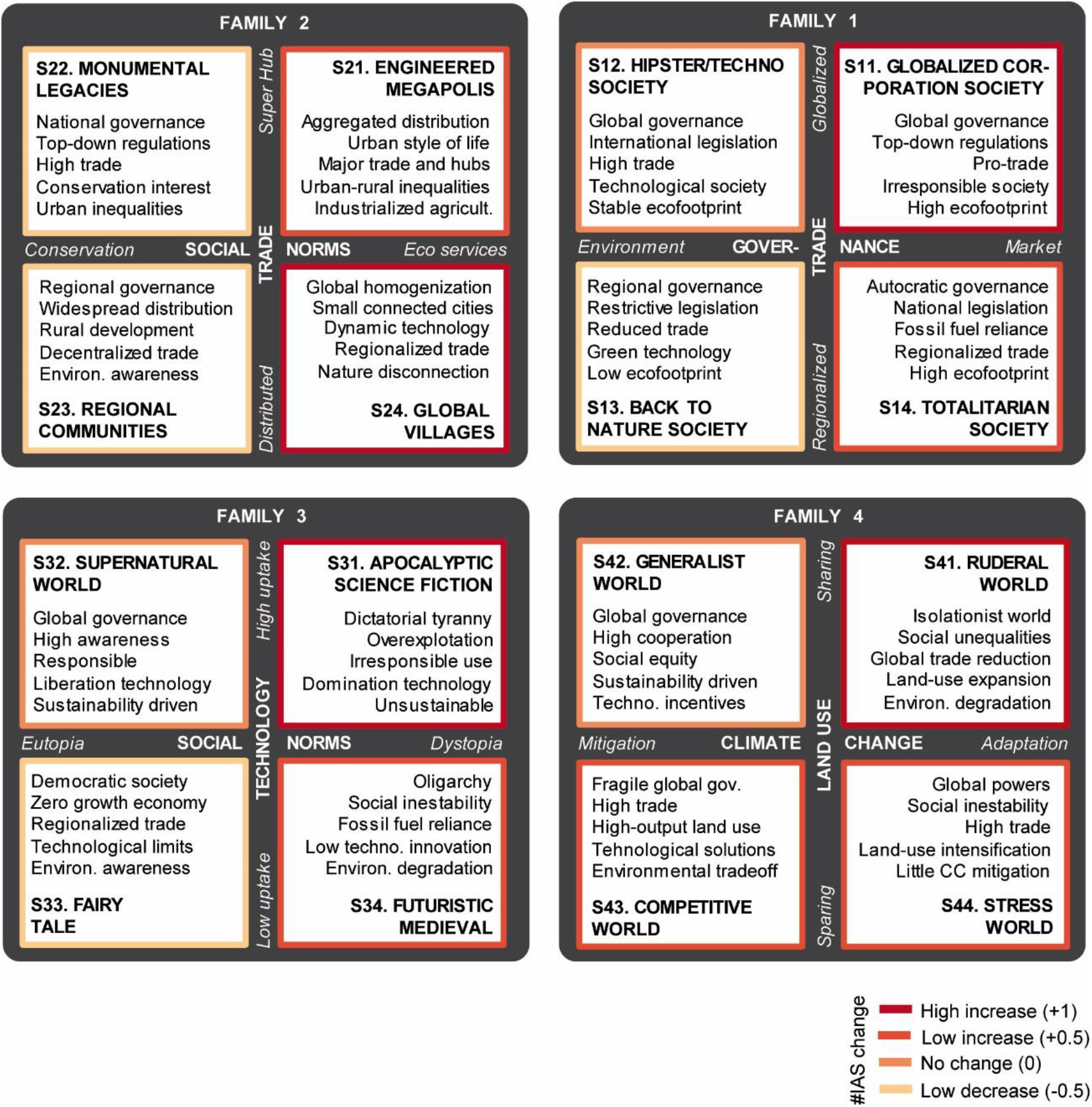
Visual summary of scenarios for biological invasions grouped by scenarios families. Scenario families are composed of two axes, corresponding to the pairs of drivers selected for building the scenarios (Fig. 2b). Red shades correspond to expert-based assessments of changes in the total number of invasive alien species under each scenarios (Fig. 4).

### Scenarios comparison

The sixteen scenarios for biological invasions identified using our novel multi-set scenario exploration span a wide range of futures, while also sharing similarities (Fig. 3; Text S1). Examining the scenarios narratives by means of qualitative content analysis, we found that the variables (i.e. terms of the content analysis) that received the highest attention across the different scenario families were those related to politics and demographic developments, as well as environmental and natural resources. Variables related to technological developments were the least considered besides its potentially enormous impact on biological invasions (Table S2). Overall, we identified 17 variables of interest to assess the similarities/differences between scenarios using a scoring rubric developed for such purpose (Table S3).

The first two components of the PCA, based on expert-based assessments of changes in these key variables (Fig. 4), explained 79% of the variance (Fig. 5). The first component (explaining 47% of variance) has positive associations with variables showing the impact of human activities on the environment and negative associations with variables related to political and social responsibility, so this component primarily measures environmental consciousness. The second component (32% variance) has negative associations with variables related to technological and trade developments, so this component measures the implementation of technology and trade advances worldwide (Fig. 5b). Scenarios ordered along these two main axes and covered the entire scenarios’ space (Fig. 5a).

**Fig. 4.**
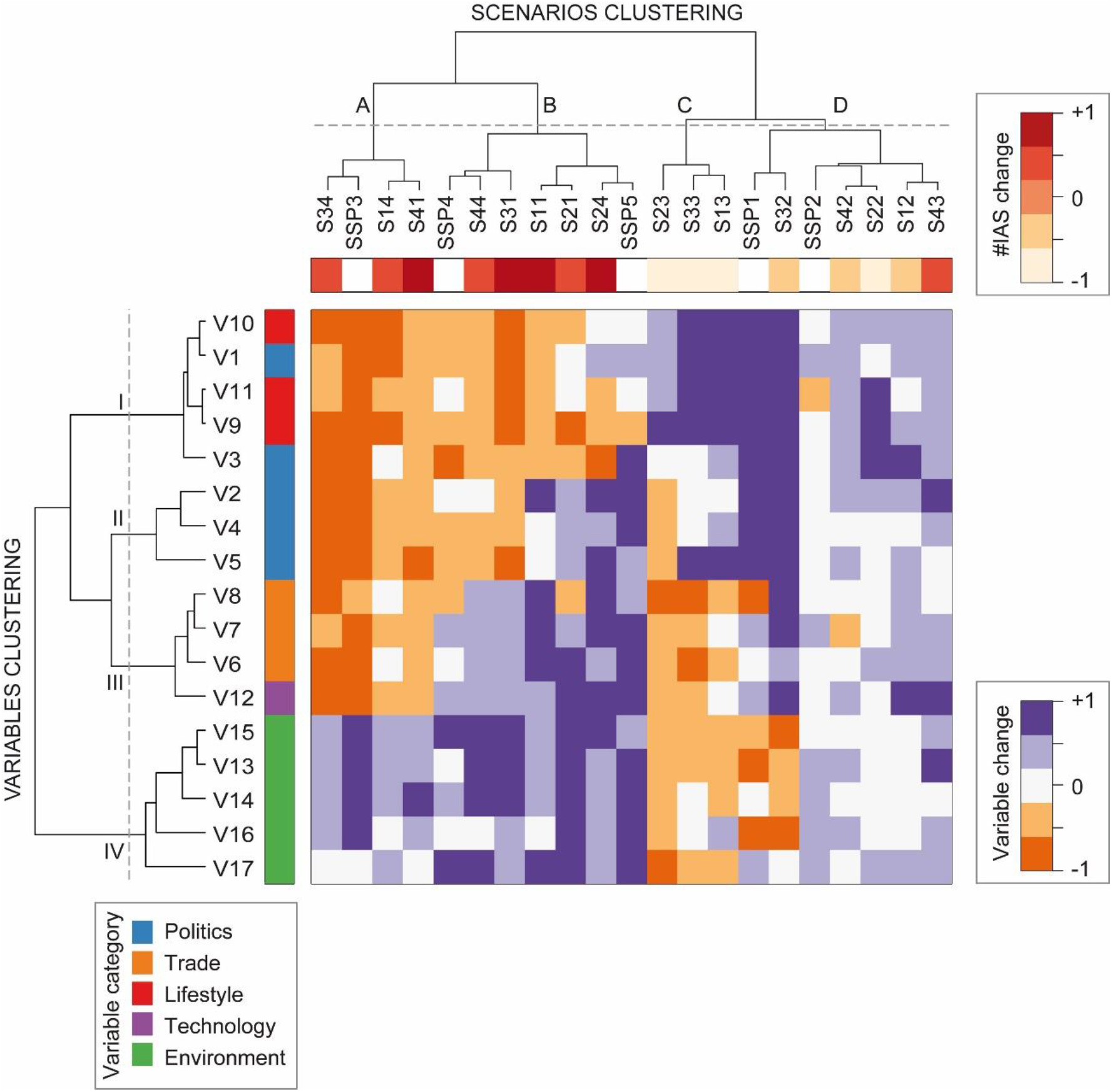
Expert-based assessments of changes in socio-ecological variables of interest characterizing the 16 scenarios for biological invasions and five Shared Socio-economic Pathways (SSP; O’Neill et al. 2017), and clustering of scenarios/variables based on these changes. Orange-blue shading intensity of the central matrix indicates the change in variables (in rows) under each scenario (in columns). The upper horizontal bar shows the change in the total number of invasive alien species, while the left vertical bar shows the five broad categories into which the 17 variables are classified. Changes in variables and the total number of invasive alien species are described in a 5-level scale from − 1 (high decrease) to +1 (high increase) (Table S3). Dendrograms represent the similarity of scenarios/variables based on hierarchical cluster analysis. Variables (coded as V#) are described in Table S3, while scenarios for biological invasions (S#) in Text S1.

**Fig. 5.**
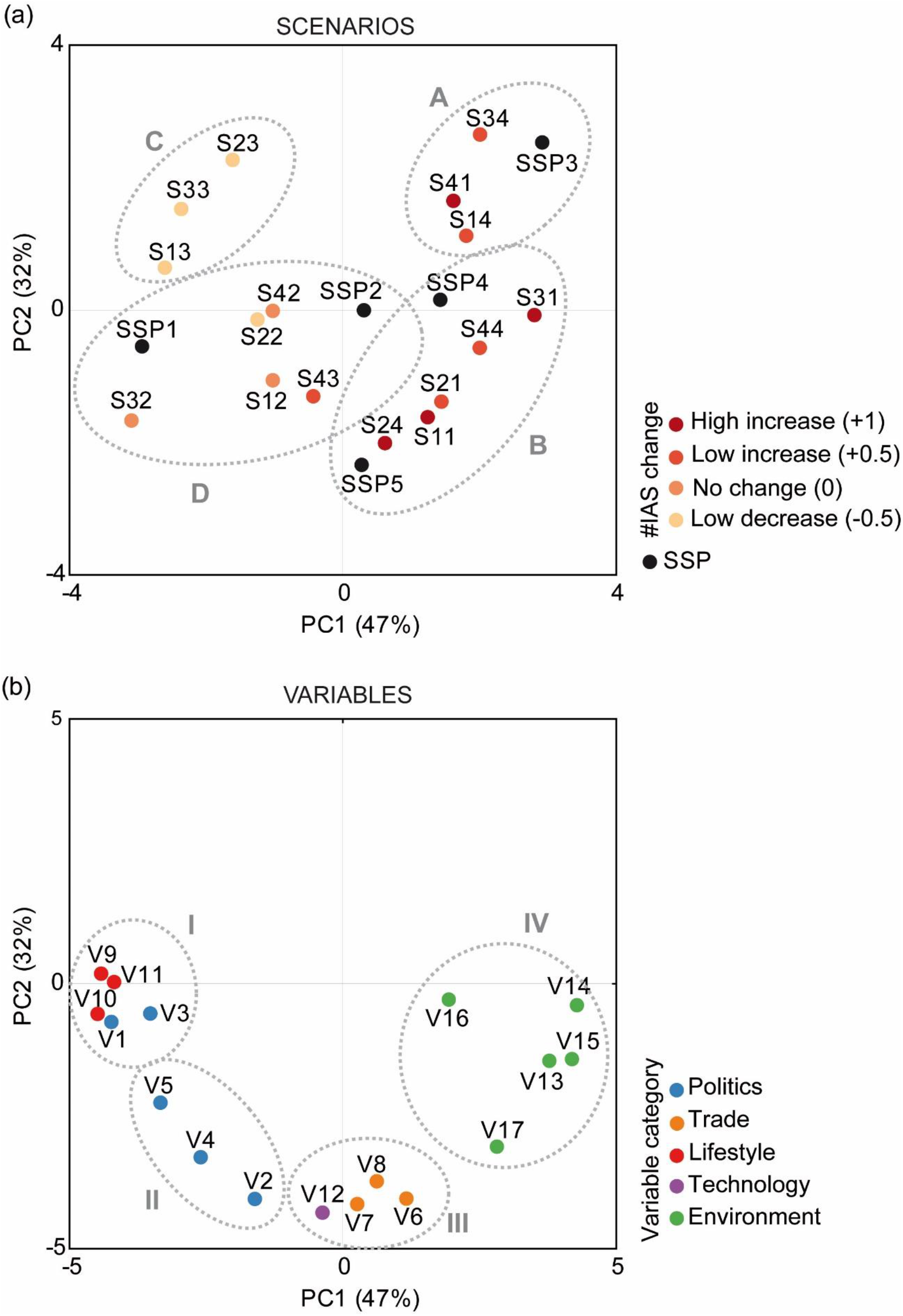
Principal component analyses on Spearman covariance matrix showing the relationships between scenarios for biological invasions and shared socio-economic pathways (SSP; (O’Neill et al. 2017). Graph (b) presents the correlations of the socio-ecological variables of interest used to characterize the scenarios (Fig. 4, central matrix) with the first two components of the principal component analysis. The dashed circles and letters/roman numerals correspond to the clusters found in Fig. 4 for both scenarios and variables, respectively. The codes for variables (V#) and scenarios for biological invasions (S#) are listed in Table S3 and Text S1, respectively.

Using the scenario coordinates in the main two axes of the PCA, we clustered the scenarios into four groups (representing the four corners of a scenarios’ space). These groups were coherent with respect to the level of biological invasions of the scenarios they included. Two groups corresponded to futures with higher level of biological invasions (A and B) and two others were characterized by lower levels of invasions (C and D) (Fig. 4, upper horizontal bar; Fig. 6). In addition to scenarios, we also clustered the variables based on how they co-varied across scenarios (Fig. 4, left vertical bar). Futures with rising numbers of IAS were associated with increasing levels of human pressure on natural environments (cluster IV), while declining trends in biological invasions tended to occur in futures with increasing levels of sustainability policies and lifestyles (cluster I) and global governance and social stability (cluster II). Increasing levels of technological development, transport, and urbanization (cluster III) were indistinctly associated with scenarios with both a high and low level of biological invasions (Fig. 4). Variables within the same cluster appear highly correlated (Fig. S2). These scenario/variable clusters are also evident from the PCA (Fig. 5).

**Fig. 6.**
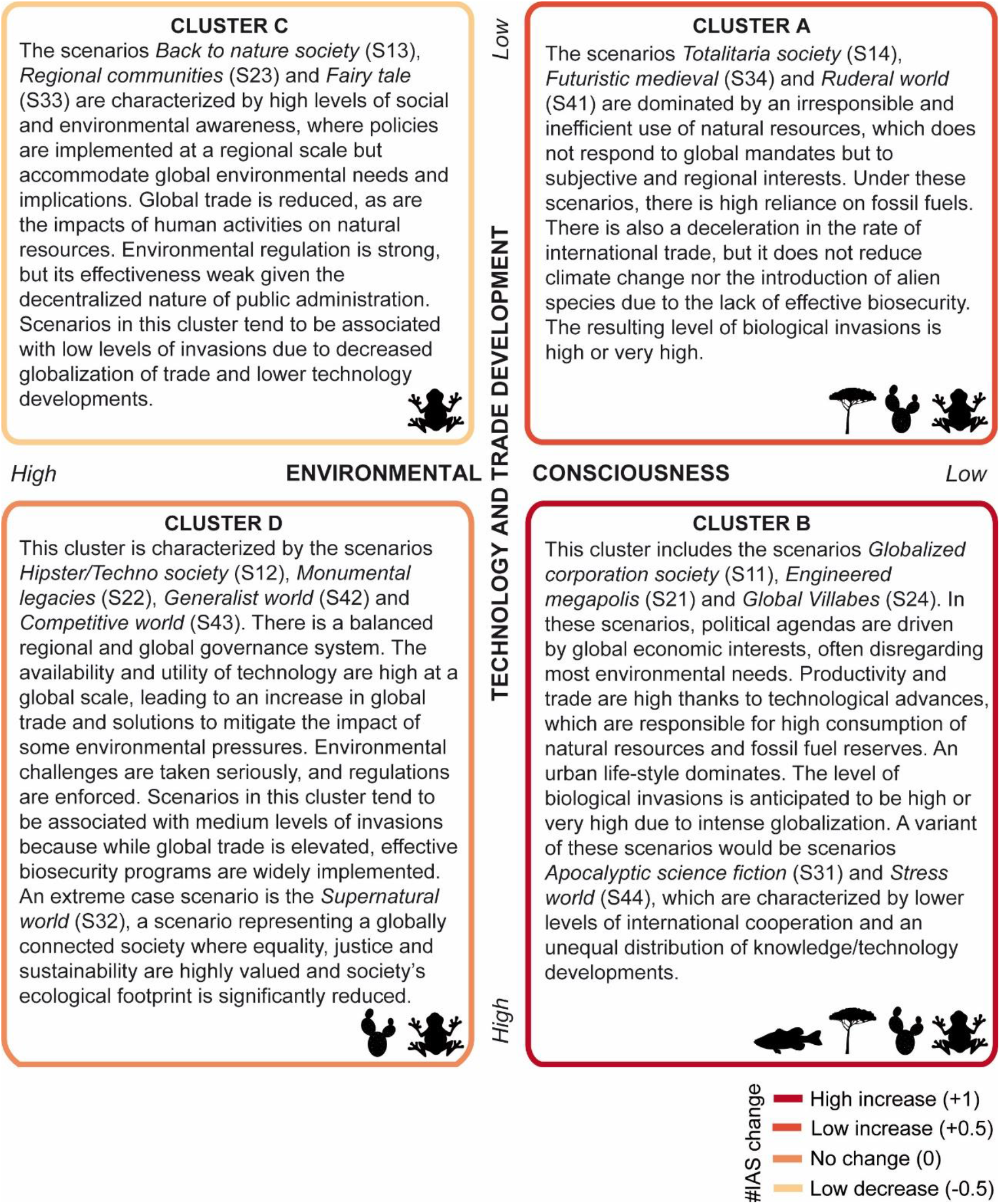
Summary of clusters of scenarios for biological invasions derived from a hierarchical cluster analysis performed on expert-based assessments of changes in socio-ecological variables of interest under the 16 scenarios (Fig. 4). The description associated to each cluster has been elaborated based on the characteristics of the scenarios included in each group (Text S1). Red shades and icons of invasive alien species are proportional to the number of invasive alien species expected for each cluster by 2050.

Some of our results do not necessarily follow the common perception that biological invasions are only associated to increasing trade and economic growth, rather we found that how society develops (e.g. economic and social development, IAS mitigation strategies) is relevant to increase or reduce the risk posed by IAS. For example, the scenarios in cluster A present high levels of biological invasions despite declining trends in trade and transport (Fig. 6). Instead, expert assessments suggest an increasing number of IAS might also result from reduced advances in socio-political developments, and sustainability policies and lifestyles, together with high levels of human modification of natural environments. While recent analyses of biological invasions have incorporated surrogates of such human modification and economic wealth, few analyses have included variables related to sustainability, such as levels of environmental awareness or development of sustainability policies (Sardain et al. 2019; Seebens et al. 2017). Our results suggest that understanding the future of biological invasions requires more interdisciplinary research that analyses how social and ecological drivers interact to shape biological invasions (Kueffer 2017; Ricciardi et al. 2017).

### Comparison with Shared Socio-Economic Pathways

Following the development of our qualitative scenarios for biological invasions, we compared them to the shared socio-economic pathways (SSPs) (O’Neill et al. 2017). Clusters A and B which have higher level of biological invasions appear fairly well represented by the SSPs. The scenarios in cluster A share features with the relatively pessimistic SSP3 (named *Regional rivalry-a rocky road*) in which the world disintegrates politically and economically into smaller regions, while most scenarios in cluster B would be more similar to the high economic growth pathway SSP5 (*Fossil-fueled development - taking the highway*) reliant on very high levels of fossil fuel use. Two scenarios in cluster B (S31 and S44) would, however, share many elements with the highly unequal SSP4 (*Inequality - a road divided*) world (Figs. 4 and 5).

Conversely, futures characterized by lower levels of IAS (C and D) are only partially captured by the SSPs. The scenarios in cluster D have been associated with the middle path SSP2 (*The middle-of-the-road*) where trends do not shift markedly from historical patterns (O’Neill et al. 2017). An extreme case scenario is S32, which shares many features with the relatively optimistic SSP1 (*Sustainability-taking the green road*) that is oriented towards sustainability. While cluster D shares features with SSPs, scenarios in cluster C describing a world characterized by regional sustainable developments that present the lowest levels of biological invasions do not. Scenarios in this cluster could be similar to an SSP1 variant with rapid shift to lower consumption lifestyles (Figs. 4 and 5).

The comparison of SSPs to our scenarios reveals that existing environmental scenarios do not represent all variables that shape biological invasions. The SSPs were created to represent different combinations of socio-economic challenges for the mitigation and adaptation to climate change (O’Neill et al. 2017), but they lack key influential biologically-oriented variables (such as biosecurity) responsible of major changes in future biodiversity. Some of these variables appear to co-vary with variables within the SSPs, but other do not (Figs. 4 and 5). We suggest that to include biological invasions in global environmental scenarios requires: (1) consideration of a broader range of positive scenarios (such as those included in cluster D) in order to capture the widest possible range of biological responses, and (2) incorporation of key drivers/variables relevant for biological invasions that are currently missing from SSPs. Futher understanding of these drivers/variables and their influence on other relevant variables is essential to enable quantitative analysis of how alternative future societal dynamics could shape biological invasions (Lenzner et al. 2019).

## Conclusions

Ecological and some economic factors have captured most of the attention of invasion science, but our expert-based analysis signals the primary role of socio-economic developments and sustainability policies and lifestyle as important drivers of biological invasions. Analyses of how trade and transport dynamics shape biological invasions have provided a starting point, but understanding the future of biological invasions requires analyzing how variables such as technological innovation, urbanization, wealth inequality, social stability, biosecurity and sustainability policies interact with one another to determine biological invasions. Realistic assessments of future biological invasions can only be achieved by considering a broader diversity of factors than are currently addressed, and doing this requires research that more broadly examines how people and societies interact with biological invasions. It is especially necessary to consider drivers and responses of biological invasions explicitly (rather than implicitly by relevant covariates) to produce meaningful results.

Our novel scenario development method has proven to be an important component in allowing us to engage with a multi-dimensional scenario space beyond any one scenario set, while our aggregation method allowed for a comparison across this larger set. We captured different potential future trajectories of biological invasions, focusing on a large variety of interacting drivers, and then grouped these multiple scenarios into four clusters (or scenario archetypes) presenting divergent futures. Future scenarios would likely be improved if they were able to better represent the global variety of experiences surrounding biological invasions, for example by ensuring that workshop participants represent the diversity of the world, balance genders, and represent different expertise. Further developments of these global scenarios and their refinement into regional or local contexts are needed to better understand the synergies between drivers/variables shaping the future of biological invasions across spatial scales. However, this work allows for the first time to establish a sound basis for global analysis of future alien trajectories and to facilitate the future quantification of the effects of biological invasions on biodiversity, livelihoods and well-being.

## Acknowledgements

This work was supported by the COST Action “Alien Challenge” [grant number TD1209]; the BiodivERsA-Belmont Forum Project “Alien Scenarios” [grant numbers AEI PCI2018-092966 (NRP) / FWF project 4011-B32 (FE, SD, GL, BL) / BMBF project 01LC1807B (JMJ)]; Deutsche Forschungsgemeinschaft [grant numbers InDyNet, JE 288/8-1; JE 288/9-1, 9-2] (JMJ); OP RDE grant EVA4.0 [grant number CZ.02.1.01/0.0/0.0/16_019/0000803] (AML); CONICYT [grant number AFB170008] (AP); UK-SCAPE programme, Natural Environment Research Council [grant number NE/R016429/1] (HER). We highly appreciate the constructive comments of an anonymous reviewer and the handling editor, Iris Bohnet.

