## Supplementary Material for "Alternative futures for global biological invasions"

**Electronic supplementary material**

**Table S1.** Participants involved in the development of scenarios for biological invasions, indicating their country and expertise. Workshop participants contributed throughout the entire process (including the presential scenario analysis meeting), while pre- and post-workshop participants contributed before and after the workshop.

| | Country | Expertise |
|---|---|---|
| **Workshop participants** | | |
| Sven Bacher | Switzerland | Invasion biology |
| Stefan Dullinger | Austria | Invasion biologist |
| Karl-Heinz Erb | Austria | Global change biology |
| Franz Essl | Austria | Invasion biology |
| Jonathan M. Jeschke | Germany | Invasion biology |
| Stelios Katsanevakis | Greece | Invasion biology |
| Ingolf Kühn | Germany | Invasion biology |
| Brend Lenzner | Austria | Invasion biology |
| Brian Leung | Canada | Invasion biology |
| Andrew M. Liebhold | USA | Invasion biology |
| Michael Obersteiner | Austria | Global change biology |
| Anibal Pauchard | Chile | Invasion biology |
| Garry D. Peterson | Sweeden | Global change biology |
| Wolfgang Rabitsch | Austria | Invasion biology |
| Nuria Roura-Pascual | Spain | Invasion biology |
| Helen E. Roy | United Kingdom | Invasion biology |
| Lucas Rutting | The Netherlands | Scenarios analysis |
| Hanno Seebens | Germany | Invasion biology |
| Marten Winter | Germany | Invasion biology |
| **Pre-/Post-workshop participants only** | | |
| Mark A. Burgman | United Kingdom | Invasion biology |
| Piero Genovesi | Italy | Invasion biology |
| Philip E. Hulme | New Zealand | Invasion biology |
| Reuben P. Keller | USA | Invasion biology |
| Guillaume Latombe | Austria | Invasion biology |
| Melodie A. McGeoch | Australia | Invasion biology |
| Gregory M. Ruiz | USA | Invasion biology |
| Riccardo Scalera | Italy | Invasion biology |
| Michael R. Springborn | USA | Environmental economy |
| Joost Vervoort | The Netherlands | Scenarios analysis |
| Betsy von Holle | USA | Invasion biology |

**Text S1.** Narratives of scenarios for biological invasions created during the scenario-based analysis, which include a description for general developments (on political and institutional developments; socio-economic and demographic developments; culture, norms and values; technological developments, science; and natural resources, ecological developments) and specific details on biological invasions. Scenarios appear grouped by scenarios families (i.e. group of four scenarios developed using the same pair of drivers).

**FAMILY 1** (by Stefan Dullinger, Andrew M. Liebhold, Wolfgang Rabitsch, Marten Winter)

Framing the drivers:

**(1) International politics, governance and legislation** (abbreviated *Governance*) – A conglomerate term of different processes undertaken by a government or organizations and influenced by external factors such as NGOs or media. We framed this driver along two opposing objectives, with ultimate goals that largely exclude each other. High focus on restrictive biosecurity policies. **A)** *Environmentally oriented governance*: processes focusing on recognizing, protecting and supporting the environment per se and/or environmental services to human well-being, underpinned by an effective biosecurity policy either through the sustainable use of natural resources or a technological solutions to environmental problems. **B)** *Market oriented governance*: policies focusing on short-term profit through abandoning or downregulating market instruments including biosecurity and disregard for environmental problems.

**(7,9) Global trade** (*Trade;* composite of drivers: (7) *economic* growth, and (9) *global trade*) – The exchange of goods at all scales from local to global. We framed this driver along two not completely opposing directions as both can be intermingled to some extent, depending on the goods. **A**) *Regionalized trade*: people exchange goods at local to regional scales. This may occur either through deliberate abstinence of consumption or through constrained options due to external factors such as societal conflicts. **B)** *Globalized trade*: people exchange goods at the global scale, with increasingly efficient transport technology. A high diversity and volume of traded goods is expected and a high level of international cooperation.

**Scenario S11: Globalized Corporation Society** (market oriented, globalized trade)

In this scenario governance is executed by one or a few global corporations. It is not a dictatorship, but prioritization of economic markets and global trade has forced jurisdiction to be largely centralized. Although national states and borders still exist, decisions and rules are enforced at large scales in a strictly top-down process. There are pro-trade regulations in place, with few, weak or abandoned regulations with regard to social and environmental protection. Ineffective biosecurity measures prevail, all leading to a high permeability of borders. The diversity and volume of traded goods is high, as is propagule pressure of IAS, transport distances are large without restrictions and overcome by expansion of existing or the creation of new pathways for introduced species. For example, global shipping volumes increase by constructing and using larger ships, new canals and waterways; unconstrained global trade of pet animals and ornamental plants increases. This leads to a globalized society with a high environmental footprint, a large middle class with a high consumption rate of natural resources, and little care for future generations. Energy production is largely carbon-based and environmental pollution very high. People prefer living in highly urbanized mega-cities and enjoy a high degree of mobility across the globe. Population growth is unabated / high and the level of cultural homogenization is also high. Most natural ecosystems are destroyed or highly degraded, biodiversity loss is very high, also due to a lack of mitigation or adaptation measures regarding climate change or land-use change, all of which further accelerate. Catastrophic events (fires, floods) are accepted as unavoidable and only post-hoc action is taken.

*For IAS*: The number of introduced and established IAS in this scenario is very high due to high propagule pressure and ineffective biosecurity measures, strongly reduced biotic resistance, and positive interactions (facilitation) with other drivers of global change. Novel communities and ecosystems dominate. Loss of biodiversity and ecosystem services is unabated and increasing. Number of IAS having negative impacts on plant, animal and human health is high, negative effects are common and response capacities are limited. No precautionary countermeasures are taken into consideration.

**Scenario S12: Hipster/Techno Society** (environmentally oriented, globalized trade)

In this scenario governance is executed by democratic societies with a balanced regional and global governance system. Life is highly and strictly regulated through international co-operation and legislation that also includes strict enforcement of highly effective biosecurity measures. The diversity and volume of traded goods is high, as is propagule pressure of IAS. Pro-trade agreements are in place, trade also increases due to global e-commerce and new trade technologies. Societies are characterized by a strong belief that technical progress can solve all current and future problems. There are high technological and educational standards with a heavy emphasis on science and technology. Economic equality is high, but there may be an aristocratic-like upper techno class. Societies have a high, but stabilized ecological footprint. There are technological solutions in place for dealing with climate change (climate engineering, renewable energy production, low-carbon technology) and land-use change (e.g. advancing use of existing agricultural land, aquaculture, GMOs). Further biodiversity loss is almost halted. Natural ecosystems are highly transformed or pristine habitats maintained under strong protection for recreation or prioritized ecosystem services. People live in large cities; there is low population growth and long life expectancies. Catastrophic events (fires, floods) are under technological control.

*For IAS*: The number of introduced IAS is still relatively high because of intensive global trade and transport, but lower than in Scenario 1 because of highly effective pre-border biosecurity efforts. The rate of IAS establishment is low, because of strong and diligent biosecurity measures, efficient risk assessments and other precautionary measures. IAS are largely restricted to cities and semi-urban environments in highly technological and artificial novel ecosystems. Facilitation of IAS through other drivers of global change is limited, but has not stopped. Further loss of the remaining biodiversity is low or stopped and loss of ecosystem services is even lower or artificially enhanced. Number of IAS having negative impacts on plant, animal and human health is low as the spread of these species is under control due to high technological response capacities.

**Scenario S13: Back to Nature Society** (environmentally oriented, regionalized trade)

In this scenario governance is executed at a decentralized, local or regional level. There are more restrictive, and less globally harmonized regulations concerning trade and the exchange of goods. Biosecurity is important, but because there is so little global trade, it is not greatly enhanced and borders have a low permeability with regard to propagule pressure. Technological advances are promoted only in relevant (sustainable) areas of research. Income disparity is low in these egalitarian societies that care for future generations. There is lower diversity in traded goods, transport distances are short, local production is implemented with regard to environmental goals and standards (e.g. organic farming, no GMOs) and propagule pressure of IAS is low. The ecological footprint of the society is small, ecosystems with long turn-around times (old growth forests, bogs) and biodiversity hotspots are secured. Pets and ornamental plants are recruited from local stocks. People live in smaller cities or rural communities, level of urbanization is low, natural resources are used effectively (e.g. renewable energy, low-carbon economy), and individual mobility is reduced. Population growth and land-use change can be high or low. Climate change has been stopped or the rate slowed down. Catastrophic events (fires, floods) are reduced by increasing resilience of the environment.

*For IAS*: The number of introduced and established IAS is very low due to regionalized trade patterns and low diversity of goods exchanged over long distances (low propagule pressure) and rigorous, though relatively simple biosecurity policies are in effect. Response capacities are well developed and sufficient to keep numbers and impacts of IAS at a low level. Facilitation of IAS through other drivers of global change is limited as these factors are halted or reduced. Loss of biodiversity and ecosystem services is low, as well as are impacts on plant, animal and human health.

**Scenario S14: Totalitarian Society** (market oriented, regionalized trade)

In this scenario there is strong, autocratic governance with a centralized jurisdiction at a national level. There are few, but very strong biosecurity measures, e.g. dedicated border control agencies, acting at invasion hubs or hotspots. Pro-market instruments are selectively downregulated to promote regionalized trade with exclusive partners within the nation and short transport distances; there is little international collaboration with widespread nationalism and xenophobia in these totalitarian societies. There is lower diversity and a reduced volume of goods concentrated on selected exchange routes among allied nations, and characteristically shortages of natural resources. Pets and ornamental plant exchange decreases. The ecological footprint of this society is high; people live in cities with a large lower class with high population growth rates, but also high mortality rates due to conflicts and reduced medical care, or isolated rural communities. Energy production is still carbon-based and environmental pollution very high. Individual mobility is low for recreation, but there is a high amount of human migration due to inequalities (economic, education, health care), including conflicts between governments and within society. Government investments into research and development as well as social sciences are low. Natural ecosystems are under high pressure, highly degraded or destroyed. Biodiversity loss is very high, nature conservation and environmental protection goals are non-existent. Climate change and land-use change accelerate, although less than in Scenario 1. Catastrophic events (fires, floods) are accepted as unavoidable and post-hoc action is taken only for privileged regions. The risks of political instability due to governmental or social conflicts are high (e.g. bioterrorism, corruption).

*For IAS*: The regionalized trade and isolation tendencies as well as the focusing of biosecurity measures on few invasion hubs or hotspots limit the number of introductions of IAS. The number of established IAS in this scenario, however, is higher than in scenarios S12 and S13 due to the high propagule pressure resulting from the market-oriented policies and less efficient response capacities after successful incursions. In other words, the barrier between introduction and establishment is the weakest of all scenarios. Facilitation of IAS through other drivers of global change is accelerating. Loss of biodiversity and ecosystem services is very high, as well as the impact on plant, animal and human health, due also to the lack of response capacities.

**FAMILY 2** (by Sven Bacher, Bernd Lenzner, Brian Leung, Garry D. Peterson, Hanno Seebens)

Framing the drivers:

**(11-13) Culture, norms and values** (*Social norms*; composite of drivers: (11) knowledge, information and education, (12) lifestyle and recreation, and (13) social norms and values) – We explored value systems, both of which were geared towards sustainability, but with different visions. **A)** *Conservation oriented*: this set of values emphasized conservation of ecosystems in their historic states, resisting global change, and the maintenance of cultural landscapes. These sets of values focus on maintaining both relatively wild and cultural landscapes. **B)** *Ecosystem services oriented*: this set of values focused on “engineering” ecosystem to enhance human wellbeing. Engineered ecosystems are a response to global change that modifies changing ecosystems, to create “novel ecosystems” that provide enhanced supplies of ecosystem services to improve human quality of life. These values consequently emphasize the urban ecosystems in which most people live.

**(7,9) Global trade** (*Trade;* composite of drivers: (7) *economic* growth, and (9) *global trade*) – We explored different shifts in global trade networks. **A)** *Super hub*: this emphasizes the concentration of global trade through key hubs. This alternative proposes that economies of scale in transport, further centralize trade. For example, use of larger, inexpensive ships result in most long-distance trade occurring through hub ports. In this world, trade is characterized by slow, cheap, long-distance, globalized and high volume transport primarily driven through a spoke and hub system through massive ports which are surrounded by large cities. **B)** *Distributed:* this emphasizes the distribution of trade away from hierarchy towards peer to peer trade. Here we assume innovations in logistics, energy, and robotics enable inexpensive, efficient local transport (e.g., advent of driverless vehicles, drones, rapid trains, new energy sources). In this transportation scenario there is less long-distance trade, but more regional and local trade. Trade is dominated by fast, dynamic, short-distance, regionalized trade in which a high volume of goods is transported through a diverse and dynamically shifting network of connections.

**Scenario S21: Engineered Megapolis** (ecosystem services oriented, super hub trading)

The “Engineered Megapolis” world is characterized by an “aggregated” distribution. The cascading effects due to larger ships is that ports (and canals) need to also become larger to accommodate them, requiring substantial investment. There is a synergy between routing shipping through major hubs, and a focus on the urban quality of life, such that more and more resources become focused on cities with the major hubs, with consequent population migration and growth within these hub “super cities”. The quality of life is generally good in super cities worldwide, with high investment within these super cities, multiculturalism, green spaces, and recreational opportunities within and close to each city area. As populations and resources become more urbanized and centralized, so too do the drivers of political power. There is a priority placed on ecosystems that provide direct ecosystem services to society within each city, which is function oriented, and often highly engineered and managed.

There are consequences to the choice to aggregate, however. There is a premium for land, and properties are expensive. While equality is generally high within super cities, slums also exist especially with new immigrants. Agriculture becomes increasingly industrialized to provide the needed food and goods to the main cities. Consequently, agricultural practices are output oriented and follow maximum yield politics, leaving a trail of abandoned and degraded land as production moves on to more fertile regions. As urban life quality improves, there is less of a need to travel outside the cities, and these degraded areas are then abandoned to become feral. The landscape will be less fragmented and will consist of three area types: urban, agricultural and abandoned. Tourism will be concentrated to some attractive regions worldwide. While migration has primarily been towards the cities, some individuals invariably remain in these rural communities, which become increasingly poor. A major source of inequity is between urban and these abandoned rural areas. Moreover, not all regions in the world prosper equally, and there is intermediate global inequality, with some regions serving as resources for others. Nonetheless, given the importance of trade and hubs, global stability is at a premium, and it is largely an integrated world. There are ramifications for technological development as well: the increasing demand for food as the population grows, will increasingly need to be grown in built environments (e.g., greenhouses) as fertile areas continue to be overused and abandoned.

*For invasive alien species (IAS)*: There is a higher potential for hitchhikers, given more trade (especially global), bigger ships, more packing material, resulting in higher propagule doses, largely localized in hubs, where they can overcome such aspects as Allee effects. Because trade is mainly from locations far away, the likelihood that the species transported are alien to the new location is high (compared to the more regional P2P trade scenarios). For marine non-indigenous species (abbreviated NIS), there is also more artificial habitat as a consequence of larger ports. However, given slow, long distance travel, transported propagules tend to be less viable (note, we’re not sure about the effect on colonization pressure, compared to P2P green scenario). As a consequence of the concentrated human activity at hubs, human impact on nature will be unequally distributed as well, with natural sites and abandoned feral sites far away from hubs being less affected and less invaded compared to super cities and their surroundings.

This is a highly controllable system, where border control can be implemented given the hub like system, and biosecurity is more effectively enforced. However, biosecurity is not given a high priority and, therefore focused only on species that are known to negatively impact human and livestock health or agriculturally relevant crops (black lists). Establishment of alien species will be mainly concentrated around hubs and will be restricted to species that can cope with the urban habitats around ports. It is a reactive system that primarily takes action after damages accrue. This system is not concerned with conservation, and ornamental, charismatic flora and fauna, and GMOs are liberally introduced. Given the high aggregation of exotic flora and fauna, extreme events (e.g., floods), although they occur rarely, could release a great number of cultured species and those accidentally introduced aliens that established around hubs. Once released, several factors are at play for subsequent spread. First, there is substantially less regional recreational travel, which reduces regional spread. However, spread between city hubs remain high. Further, regions outside cities have less resources, fewer people and less surveillance. Taken together with land abandonment, high biodiversity loss, this disturbed landscape provides fertile ground for NIS to survive and spread. Given the highly intensive industrial farms, there is a substantial risk of food insecurity due to invaders.

**Scenario S22: Monumental legacies** (conservation oriented, super hub trading)

The "Monumental legacies" world largely follows and extends today's conservation approaches. It continues the trend towards larger ships, and towards conserving land areas (e.g., protected areas), and the character of rural communities, for future generations. Here there is substantial interest in preventing NIS and a concern about biodiversity, beyond its direct utility for society. While there is a tendency towards bigger ships, and the consequent larger ports and canals needed to support them, emerging trade-hub cities, and aggregated resources, it does not reach the same level as "Engineered Megapolis", since there is an explicit desire to focus on the entire landscape. Nonetheless, there is national, centralized top-down regulations, national parks and conservation initiatives. There are strong environmental regulations and legal enforcement. The landscape is maintained for the purpose of conservation (and recreation), hence fragmentation is lower than today, but higher compared to "Engineered Megapolis". A premium is placed on biodiversity, maintaining genetic pools, and having adaptive landscapes. Environmental awareness in the public is very high and results in the development of more small-scale initiatives that increase biodiversity (i.e. seed trade/markets, conservation of old lineages). Nonetheless, while more resilient over the long-term, food production is a challenge, with less absolute production efficiency afforded by industrialized farming approaches.

The environment is as important as society, and thus, there are sometimes forced human relocations, and human migration to cities. In these cases, there is uncontrolled urban growth, with problems of slums. This results in relatively high urban inequality, but much less rural versus urban inequality, since resources are invested widely to conserve the natural and cultural landscape, and land abandonment is slow. At the global level, inequality is similar to current levels.

*For IAS*: Given the strong environmental ethos, biosecurity and NIS risk screening are strong, the central shipping routes are well regulated, and prevention is effective. Moreover, while container ships are larger resulting in more hitchhikers, the distances transported are longer and slower, and thus, the overall propagule pressure is lower. Genetically modified organisms, assisted migration, and other purposeful introductions are relatively rare and heavily regulated, and conservation of old crop lineages and their genetic diversity is supported. However, given the large national parks system and recreational opportunities outside the city, there is high regional traffic, and therefore a high propensity to spread NIS. Once they spread, though, the high biodiversity leads to higher biotic resistance, and a higher probability of detection as people are more widely distributed and generally aware of environmental threats. However, societies are economically poorer, and thus there are fewer resources available to manage NIS.

**Scenario S23: Regional communities** (conservation oriented, distributed trading)

This is a very socially/environmentally aware world, with decentralized regulations and a focus on conservation of historical environmental and cultural states. The cities are smaller, there is only minor land abandonment due to environmental degradation, and this world has the highest rural development and widest distribution of people, given the decentralized and flexible nature of trade. Basically, people can obtain whatever they need, regardless of where they are, as long as it is not considered a threat to the environment. Therefore, there is much less incentive to aggregate in super cities. Due to the wide geographical distribution of the population, there is also a wider diversity of cultural norms, high regional pride, and generally low migration. There is federalism, with division of power to regional levels of government. There is also a deep respect for nature. Here, social monitoring is as important as explicit regulation. Food production remains local and more traditional, with less emphasis on scientific advances.

In this world, while environmental regulation is strong, it is less effective given the decentralized nature of trade and administration. Moreover, there are larger inequalities between regions, coordination is limited, and resources to respond to environmental threats are heterogeneous. Finally, given the higher regional pride, there is also greater regional conflict, which causes periodic migrations due to refugees.

*For IAS*: Prevention is deemed important, but given the diversity of trade methods, it is less effective. Overall, there is less long-distance trade, resulting in lower propagule and colonization pressure. However, there is more regional trade, regional travel for recreation. Consequently, while it is less likely that a new NIS establishes, if one does, it can quickly spread due to high regional transport. While surveillance is high, given the interest in conservation and relatively homogeneous population distribution, there is a lower ability to form a coordinated response across regions, and lower incentive to conduct costly management to benefit neighboring regions. In other words, this system is prone to suffer from the “tragedy of the commons”. Nonetheless, given the high biodiversity, there is high biotic resistance, and the landscape is heterogeneous and patchy, thus making the establishment and natural spread of new alien species more difficult. These act to reduce establishment, slow spread, and moderate impact – while there is a higher probability of an outbreak, the effects are more localized and are not catastrophic compared to homogeneous engineered landscapes.

**Scenario S24: Global villages** (ecosystem services oriented, distributed trading)

Global villages consist of smaller but highly connected cities, with cheap autonomous transport. Here, traffic is as likely to occur between cities in different parts of a country as in neighboring regions. Cities are more geared towards subcultures (e.g., art, religion) and specialization than offering all amenities to every subgroup. People feel like global citizens and are not attached to a specific country. The population is highly mobile, both professionally and for recreation (like academia). There is high turnover of people within any given city; cosmopolitan but not stationary. As such, there is global homogenization, but regional diversity. Given the dynamic, flexible nature of trade, people can get whatever they want wherever they live. Populations are still centered in urban environments however, and there is less interest in areas outside cities. There is a premium placed on ecosystem services directly related to human well-being, and hence novel ecosystems are widespread and diverse, with each city deciding their own needs. Technological advances are dynamic, often comprised of small start-up companies that react flexibly to new demands. These new technologies allow increased food yields, and best practices spread quickly across cities and nations. Population growth is slower, due to the non-sessile, highly educated lifestyles, which do not encourage many offspring.

While cities are still green, with city parks, and good ecosystem services, there is a general disconnection from “nature”, few “pristine” areas, and less interest in biodiversity as an intrinsic good. Regulation is geared more towards enabling travel, ecosystem services and distributed trade. Environmental regulation is less coordinated and less effective, and in many ways of less interest, except where it directly affects societal welfare. However, given the higher degree of mobility, the smaller cities, and the less aggregated nature of resources, there is a greater ability to adapt to changing conditions (e.g., climate change) due to the high creativity of society or the willingness to geographically relocate residences, but there is a higher chance of any given city being affected (compared to super cities with high investments in infrastructure).

*For IAS*: Biosecurity regulation is weak, because of low interest in prevention, low coordination, and low effectiveness (distributed network). There is less long-distance trade in terms of volume (compared to super hubs), but more trade events. Propagule pressure is more distributed (i.e., lower propagule pressure at more sites) and less concentrated to single sites like in the Megapolis scenario. However, there are many novel trade routes, which increases the number of species transported (colonization pressure). Moreover, trade is more regionalized and distances for commodity transports are on average shorter and faster, resulting in a greater number of viable propagules. On the other hand, major pathways of introduction (e.g., solid wood packing material, hitchhiking in containers) are less of an issue in this world, since weight is at a premium given the smaller modes of transport (e.g., drones). Given the interest in novel ecosystems, GMOs, exotic agriculture/aquaculture and pets, assisted colonization, are the norm, and differ between cities. These can easily spread from source locations given lower coordination, and there are continuously new invasions. The distribution of people also means that there is a higher ability to detect NIS, although this is not a priority in this world (low awareness), except for those NIS that start causing high damage to human health and goods. However, cumulative impacts of a given invasive alien species are generally lower after spreading, because NIS encounter a highly heterogeneous landscape, due to the diversity of novel ecosystems, and thus widespread devastation is less likely.

**FAMILY 3** (by Jonathan M. Jeschke, Stelios Katsanevakis, Núria Roura-Pascual, Helen E. Roy)

Framing the drivers:

**(11-13) Culture, norms and values** (*Social norms*; composite of drivers: (11) knowledge, information and education, (12) lifestyle and recreation, and (13) social norms and values) – We explored a gradient from an eutopian to a dystopian society with regards to environmental changes. **A)** *Proactive society*: well-informed and altruistic society, where human-human and human-environment relations are constructed on the basis of equality, responsibility and justice. **B)** *Reactive society*: ill-informed, selfish and totalitarian society, unable to construct positive and equalitarian relations for the benefit of its inhabitants and the environment.

**(14) Technology and innovation** (*Technology*) – Gradient from a technophilic to technophobic society. **A)** *High uptake*: technophilic society characterized by a strong transfer and adoption of new technological innovations. **B)** *Low uptake*: technophobic society where there is a low development, transfer and adoption of advanced technology solutions.

**Scenario S31: Apocalyptic Science Fiction** (reactive society, high technology uptake)

Autocratic, dictatorial tyranny with profit-driven hierarchical and compartmentalized societies lead to increased inequality and corruption. There is a lack of international cooperation with a high level of social and transboundary conflicts (often leading to terroristic attacks and wars) with associated loss to human populations, but coupled with high population growth. Economic growth is likely but wealth is distributed disproportionately among individuals; economic collapse is likely at a tipping point of resource consumption. A few countries and multinational companies control a large percentage of the global wealth. The majority of the human population has almost no direct experience and interaction with nature, with increasing interest for simulated experiences. Unsustainable use of the natural capital; energy, resources and water monopolised by a few and used inefficiently. Overexploitation of natural resources for short-term profit, which leads to environmental degradation due to pollution, habitat destruction, increased urbanisation, etc. Extreme biodiversity loss. Artificial intelligence extending beyond human intelligence. High global connectivity and technological/knowledge advances but for the benefit of a few. Inequalities in access to data, information and tools. Innovative transport systems but lots of junk/useless technology. The perpetuation of this situation, over the long term, leads to the use of artificial ecosystem-services and the exploitation of other planets.

*For IAS*: Technological developments in transport increases the scale and volume of international trade, but the disinterest of governments in biosecurity and the disinformation of people on environmental problems exacerbate the movement and establishment of invasive alien species. This is a consumerist society, where IAS are only considered problematic when they are responsible for major negative economic impacts. Tourism and human migrations due to political instability increase the movement of exotic species. The growth of population, the expansion of urban areas and artificial habitats, the increasing pressure on non-urbanized areas for food consumption and energy production, and the large-scale environmental degradation will reduce the resilience of ecosystems, cause an increased rate of biodiversity loss, and create new opportunities for alien establishment and spread.

**Scenario S32: Supernatural world** (proactive society, high technology uptake)

Democratic multicultural societies living in peace; governments, international organisations and the private sector are operating cooperatively for the benefit and well-being of society and the environment. Governance is shifted towards direct democracy, where technology facilitates participation. Highly resilient and responsible societies in which equality, justice and sustainability are highly valued. A connected and progressive world. Individuals have free time to participate in decision making and proactively respond to socio-ecological issues. Sustainable development (including population and economic growth) is an overarching principle; natural capital and ecosystem services are valuated (and thus a cost associated to their use) and conserved. Biosecurity is given a high priority on the political agenda. Technology is developed and used for the benefit of nature and societies, including technological solutions for environmental challenges (energy and transport). There is an overarching drive for useful and accessible technology, knowledge and information.

*For IAS*: Technological advances in transport increase the scale, volume and efficiency of international trade. Coupled with increasing levels of global connectivity and economic growth, it is expected to increase the number of alien species distributed and established around the world. However, biosecurity innovations (both in detection and control of alien species), the political will to support management actions to mitigate the impacts of invasive alien species, and the increasing public awareness of and involvement in biodiversity conservation will reduce the introduction rate of alien species and particularly their impacts on biodiversity, human health and ecosystem services. In other words, alien species with no or even positive impacts, which can be identified due to the high level of knowledge and technological advancements, will be allowed to establish or actively promoted, respectively, whereas the introduction of alien species with negative impacts will be minimized, and those already established will be controlled or eradicated more efficiently. Innovative technological solutions (in transport, construction and agriculture) will make urban areas environmentally friendly and reduce the current impacts of land-use degradation and fragmentation.

**Scenario S33: Fairy tale** (proactive society, low technology uptake)

Highly democratic, self-sufficient, participatory and egalitarian society, but with a poor technological capacity although resilient due to their organizational capacity. High interest for biosecurity. Basic income provided by the government. People think globally and act locally. Politics and society are aware of their social, economic, technological and environmental limitations and challenges, but some goals maybe misguided due to a lack of technology. Decelerated society whereby population growth is limited (at a lower level than today) and zero economic growth or degrowth. People tend to live simple lives; consumption and food supply remain stable at a lower level than today. Decline of international trade and preference for local products, with increase in cultivation of local varieties and an increase in organic farming. Cottage/market gardening dominates; some but disjointed understanding of natural capital. Families try to produce part of their food (small gardens, domestic animals, ponds, beehives, etc.). Good well-being and time available for higher arts (art, literature, music, lyrics, etc.) as well as for altruistic actions. Bioconservatism and biosecurity are considered a high priority in both the society and politics. Because of limited capacity to generate more environmentally friendly technology, there is a high reliance on existing technology and natural resources. Relatively low level of technology and low or no motivation for innovation, although there is equal and high access to knowledge and education. Climate change has increased, but has not reached the highest projections because of a non-growing population and economy. Biodiversity is valued and the environment preserved; intermediate level of environmental degradation.

*For IAS*: The decrement of international trade, as a consequence of low technological developments and economic growth, reduces the transportation of alien species over long distances. Local connectivity strengthens and unaided dispersal becomes the main pathway of alien species' dispersal. The consciousness of people in environmental challenges and their proactive participation in adopting biosecurity measures, promoted by policies for mitigation, lowers the rate of new introductions and the impact of invasive alien species, although the lack of innovative technological solutions will not allow very efficient control and eradication of already established invasive alien species. The effect of tourism and social conflicts is minimal, but the higher pressure on non-urban areas (because of people living in their own houses in the countryside and having their own gardens) increases land-use fragmentation and therefore affects IAS in contrasting ways (both increasing propagule pressure, but also reducing connectivity for dispersion).

**Scenario S34: Futuristic Medieval** (reactive society, low technology uptake)

Oligarchy, dictatorial despotism with politically fragmented and hierarchical societies governed by corrupted politics. Lack of global perspectives and little interest in the long-term benefits for societies and the environment. Political decision-making is based on the opinions and beliefs of a few and is not evidence-based. There is high political and social instability. Economic inequalities lead to profound poverty; most people are poor, a few are rich. There are frequent boom-and-bust economic cycles. High human mortality and high disease prevalence creates a survival of the fittest atmosphere. Human population density is intermediate and there are low technological and knowledge advancements and innovation. There is inadequate funding for research leading to technological stagnation and lack of innovation. Poor knowledge transfer and dissemination, which leads to an inefficient use of available technology and therefore a high dependency on natural resources/fossil fuels. Lack of awareness of societal and environmental challenges ahead; policies and politicians do not have the capacity to regulate or enforce societal/environmental changes. Unsustainable development, including wasteful energy consumption and unsustainable exploitation of natural resources, leads to: intensification of climate change due to increased emissions, high pollution, intensification of land-use change (to cover food demands), environmental degradation and a generalized loss of biodiversity and ecosystem services.

*For IAS*: The deceleration in the rate of international trade, as a consequence of low technological developments and stagnation of economy, will reduce but not stop the transportation of alien species over long distances. The regional and local network of transportation and the extension of urban areas will continue as present, but the incapacity of society to minimize (or even contain) the impacts of alien species, the absence of policies for mitigation of such impacts, and the disinterest in biosecurity will have negative repercussions in both environment and human well-being. The increasing pressure of society on natural resources, due to social instability and inequalities, will create new opportunities for the establishment of alien species. Changes in land use and climate will intensify and will exacerbate the number of alien and invasive alien species as well as their impacts.

**FAMILY 4** (by Franz Essl, Karl-Heinz Erb, Ingolf Kühn, Anibal Pauchard)

Framing the drivers:

**(16) Climate change** (*Climate change*) - Reactive versus proactive climate policy. We explored the approach towards climate change, one of which was proactive (i.e. strong CC mitigation), and one was reactive (i.e. focus on adaptation to experienced CC impacts). **A)** *Mitigation oriented*: climate change mitigation was one of the overarching goals of societies and policies, and competing, non-vital goals were considered of secondary importance. **B)** *Adaptation oriented*: climate change mitigation failed (due to lack of will and collaboration), and thus societies implemented adaptation measures reactively to perceived impacts. However, with increasing severity, adaptation measures became less effective.

**(21) Land use change/development** (*Land use*) - Spatial distribution and intensity of land use. **A)** *Land sparing*: concentrating land-use on highly productive regions combined with highly intensive land-use that is based on the adoption of new technologies, high energy and material input land use. In this scenario, global cooperation is highly relevant to ensure that centres of food production and consumption meet. Land freed from production needs is used for different purposes. **B)** *Land sharing*: this scenario is based on a globally more dispersed land-use, which is less focused on achieving highest output by virtually any available means. It is consistent with small-scale farming, including lower input land-use types. On average, lower yields have to be compensated by larger areas under cultivation that may result in higher land competition with natural ecosystems.

**Scenario S41: Ruderal World** (adaptation oriented, land sharing)

This world opts for a moderate transformation of current land use with strong elements of small-scale and (relative) low input farming (with medium gains in efficiency), high demand for conversion of additional land to be used for agriculture, while taking a re-active approach to climate change (lack of collaboration and will). Heterogeneous agricultural landscapes with many farmers are prevalent, which initially are resilient to some degree to external pressures. Pressure on remaining unused ecosystems is high, and substantially rising over time as CC impacts on food production increase. Landscape connectivity will be conserved, but will degrade later with increasing pressures. This scenario is consistent with an isolationistic world (that may include regional nuclei of strong collaboration, e.g. North America, Europe - EU), poor governance of natural resources, and rising inter-country inequalities, and rising risks of conflict. Food security is regionally organized, regional crop failures are more difficult to address. Technological progress is relatively low. Trade intensity may be relatively low, food is largely distributed regionally, and later, when food security declines, food prices rise, countries producing food surpluses limit the export of food. Changes in diets and/or malnutrition become regionally endemic, strong migration pressure from disadvantaged to advantaged countries.

- Political and institutional developments: Rising pressures over time increasingly leads to an isolationist world (incl. the dissolution of the EU), little collaboration and international distrust, and risks of war and conflict. However, this may include networks of regional collaboration between countries with shared interests.
- Socio-economic and demographic developments: Unequal socio-economic growth, increasing and high risks of failed states, causing large global migration pressure.
- Culture, norms and value: Dominance of self-focused strategies, including the exclusion of migrants, and the emergence of unseen levels of violence and conflict. Long-term environmental issues are considered irrelevant.
- Technological developments, science: Scientific solutions and their societal uptake are driven towards short- and medium-term challenges, while long-term questions are insufficiently addressed. Innovation in general is low to moderate.
- Natural resources, ecological developments: In short/medium term moderate impact on biodiversity, but over time strong degradation of biodiversity due to aggressive expansion of land use to secure food production.

*For IAS*: This scenario that is characterized by a preference of regional political solutions, reduced levels of international trade, severe climate and a land use changes combined with moderately intensive will lead to a sharp increase in alien species. While global trade is becoming less relevant in driving invasions, land use changes will lead to a massive decline of natural habitats in many regions, and open up areas for biological invasions. Later on, the increasing severity of climate change impacts will further enhance the spread of alien species. As societies become increasingly stressed with increasing climate change impacts, and international cooperation is limited, the management of alien species that not directly threaten human livelihoods is increasingly becoming less relevant, and many alien species expand rather unchecked. In contrast, alien species that are detrimental to human health, forestry or agriculture are strongly managed, although efficiency of measures is hampered by a lack of international collaboration.

**Scenario S42: Generalist World** (mitigation oriented, land sharing)

This world is driven by the joint commitment of limiting the pace of climate change and by relying on flexible and adaptive cultivation systems. The global lands will contribute substantially to energy generation, carbon extraction (biofuel) and sequestration. Farming will be done mostly by small-scale farming, that apply new techniques, breeds etc. for resource-efficient farming. Integrated, multipurpose strategies will be important in agriculture and forestry. This will sharply rise the need for land (low yields), but at low-to-medium intensities, and will require an integrated cooperative world with good governance to avoid mal-mitigation (e.g. conversion of high-carbon ecosystems), while little productive lands are set aside for biofuel production. This world is crucially dependent on relatively high prices of agricultural produce, changes in diet (reduced meat production), and the expansion of low energy input – high output land use (intensive organic agriculture, permaculture, agroforestry). Climate change mitigation is of prime importance, and introduced species are widely used for afforestation and biofuel production

- Political and institutional developments: Strong international institutions and governance, with strong regional bodies (e.g. EU)
- Socio-economic and demographic developments: Socio-economic disparities decline (e.g. many people will be able to work in the agriculture sector), and socio-economic development is likely to be relatively evenly distributed.
- Culture, norms and value: Long-term environmental problems are seriously addressed, and the implications of these challenges are changing societal norms. However, CC mitigation is considered of prime importance, and introduced species are widely used for afforestation and biofuel production.
- Technological developments, science: Strong incentives for innovation, e.g. into land use and resource efficiency
- Natural resources, ecological developments: Ecosystem integrity of remaining natural ecosystems is largely conserved (with the exception of e.g. grasslands or degraded lands, that are afforested on a large scale), but large areas used for plantations/biofuel production etc. are highly invaded.

*For IAS*: This scenario reflects a world that is strongly committed to combating climate change and secure food production via adaptive agriculture embedded in a collaborative world. The number of alien species will continue to increase driven by the joint effects of substantial further habitat conversion, moderate climate change, the use of land for biofuel production (often based on introduced species), and a decline in the recognition of biological invasions as causing detrimental impacts when alien species provide other benefits to society (e.g. biofuel). However, the increase in alien species will be rather moderate, and it will be rather equally distributed across the globe (with some hotspots in regions that will experience substantial conversion of rather pristine habitats into intensively used ones).

**Scenario S43: Competitive World** (mitigation oriented, land sparing)

This world is characterized by opting for high-output agriculture concentrated in productive regions, while sparing land for CC mitigation (biofuel, etc.; halting high-carbon ecosystem losses) is considered equally important. Farming and biofuel production will be partly dominated by businesses, but will also involve small-scale farmers that use their land intensively (incl. permaculture, intensive agriculture). This world is shaped by a strong divergence in land uses. Land use conflicts require strong governance. Food availability is low, thus distributional aspects are key, fostering global cooperation. The agricultural and political system is vulnerable to shocks (e.g. threatening food security) that may transform the political cooperative architecture into characterized by inequalities and domination. This scenario is crucially dependent on high-output land use, high levels of local production of low-carbon energy, changes in diets, and strong carbon and agricultural price signals. Global trade will be highly relevant to distribute agricultural produce. Alien species will be widely accepted for plantations, while those being pests or human-health relevant will be strongly managed.

- Political and institutional developments: Strong international collaborations, but at risk of being transformed into an unequal one by external shocks (e.g. food crises).
- Socio-economic and demographic developments: Techno- and efficiency-driven land use solutions with high level of innovation of land-use systems and techniques. Number of people working the agricultural sector will decline moderately.
- Culture, norms and value: Long-term environmental challenges are taken seriously, and have transformative implications on values and behaviour.
- Technological developments, science: Strong focus on innovation and land-use efficiency gains.
- Natural resources, ecological developments: Conversion of high carbon ecosystems is largely halted. In less productive or degraded regions, CC mitigation land uses dominate, while in productive regions, intensive production of agricultural products dominates.

*For IAS*: In this scenario, the world will be characterized by high trade intensity, regions of highly different land-use intensities including areas of non-managed pristine environments, managed set-aside ecosystems in less productive regions used for biofuel production etc., and intensively used productive regions and moderate climate change. The number of alien species will rise rather sharply, but unequally in different regions with land use and trade being more relevant drivers than climate change. Alien species will only increase modestly in regions that are spared from land use, while habitats that set aside for e.g. biofuel production will experience rather high levels of alien species, although there will be some management. In intensively used agricultural habitats, the intensification of land-use and the introduction of new breeds and varieties will create favourable conditions for the rapid spread of alien species including pest species with high impacts. However, international collaboration, strong alien species policies and rather ambitious management will reduce the impacts of these species there rather strongly.

**Scenario S44: Stress World** (adaptation oriented, land sparing)

This storyline represents a world where society has strongly responded towards climate change, that has opted for intensifying land-use by virtually any means in agriculturally productive regions by taking advantage of natural conditions and resources at the large scale as well as technological solutions (monocultures, irrigation, pesticides, fertilizers, glasshouses, vertical farming), while land use is being abandoned in less favorable regions. Land conversion is moderate, wilderness areas return. Agriculture is based on high-input farming, with a strong drive towards plant breeding incl. GMOs (drought resistance, warmer climate), large businesses dominate the agricultural system while small-scale/ subsistence farming is being largely abandoned, and organic farming is of limited importance. This world has opted for little climate change mitigation, and thus the pace of climate change is high, increasingly stressing food security and ecosystems, and transforming the initially cooperative world to an unequal one dominated by powerful nations. Land that has been abandoned is often planted with alien plants, and combined with climate change induced stresses; these areas are characterized by high levels of invasions. Under increasing stress, norms and attitudes shift away from those considering conservation and invasion relevant (except for pests).

- Political and institutional developments: Strong international collaboration for distributing food produced that develops into a system dominated by the big global powers under increasing pressure on food security. Initially based on a political architecture of trust and cooperation. But political instability rises sharply over time, driven by shifts in societal values towards food security that lead to a politically and economically unequal world dominated by the big nations (land grabbing, imposing favourable terms of trade), while invasions are considered less relevant.
- Socio-economic and demographic developments: Few farmers left due to high pressure on subsistence and small-scale farmers, agro-companies become global key players in land use, socio-economic instability increases towards 2050 due to increasingly severe extreme climate events.
- Culture, norms and value: Sustainability and conservation are considered of little relevance, the societal focus is on short term measures and impacts.
- Technological developments, science: Focus on large-scale technological land-use solutions.
- Natural resources, ecological developments: Large negative impact on resources, biodiversity and ecosystem stability, in particular in the centres of food production.

*For IAS*: The high intensity of trade and transport that characterizes this scenario combined with rapid climate change, and intensive land-use in productive regions will first lead to a rather moderate increase in the number of alien species that will rise more pronouncedly later on. However, there will be substantial disparities in the increase of biological invasions: regions that are set aside from agricultural use and often planted with introduced trees, become heavily invaded, and IAS management is relatively modest. Less productive or agriculturally little used regions are initially little invaded, but with increasing climate change and ecosystem deterioration, alien species expand substantially later on. Finally, in intensively used regions dominated by agriculture biological invasions are becoming severe, although at least on productive lands partly managed.

**Table S2.** Results of the coding process performed on the scenario narratives for biological invasions (Text S1) to identify the variables shared by most families of scenarios (F1-F4). Variables appear grouped in different categories, as well as differentiated depending on whether they appear into the general description of the scenarios or in those dedicated to invasive alien species. Only variables shared by three or four families of scenarios (indicated in column **Σ**) were considered for characterizing scenarios. The last column (ID) indicates the correspondence with variables listed in Table S3.

| | F1 | F2 | F3 | F4 | Σ | ID |
|---|---|---|---|---|---|---|
| **i. Politics and demographics** | | | | | | |
| General context | | | | | | |
| Biosecurity | | x | x | | 2 | |
| Governance | x | x | x | x | 4 | V1 |
| Governance scale | | x | x | | 2 | |
| Inequalities, mainly economic but also to information (global/regional) | x | x | x | x | 4 | V4 |
| International collaboration | | x | x | x | 3 | V2 |
| Population growth (mortality) | x | x | | | 2 | |
| Population migrations | x | x | | x | 3 | V5 |
| Priorities (socio-political) | x | x | x | x | 4 | V2 |
| Reactive/proactive system (time availability) | | | x | x | 2 | |
| Regulations (environment/trade) and enforcement | x | x | | | 2 | |
| Social stability (conflicts) | x | x | | x | 3 | V5 |
| Invasion context | | | | | | |
| Biosecurity (effectiveness, prevention, border control, etc.) | x | x | x | | 3 | V3 |
| Human migrations | x | | x | | 2 | |
| Reactive/proactive system | x | | x | | 2 | |
| Responses/measures (coordination, precautionary) | x | x | | | 2 | |
| **ii. Economy and trade** | | | | | | |
| General context | | | | | | |
| Resources (funding) | x | | x | | 2 | |
| Trade (volume, diversity) | | x | | x | 2 | |
| Trade patterns (long- vs. short-distance) | x | x | | | 2 | |
| Transport/mobility/connectivity | x | x | x | | 3 | V7,V8 |
| Invasion context | | | | | | |
| Connectivity | | | x | | 1 | |
| Economic growth | | | x | | 1 | |
| Trade (packing material) | x | | x | | 2 | |
| Trade (ships, trade) | x | | x | x | 3 | V6,V7 |
| Urban-Rural distribution | | | x | | 1 | |
| **iii. Lifestyle and values** | | | | | | |
| General context | | | | | | |
| Care for future generations | | x | | | 1 | |
| Changes in diet | | | | x | 1 | |
| Connection to nature | x | | x | | 2 | |
| Culture (homogenization?) | x | x | | | 2 | |
| Education standards | | x | | | 1 | |
| Quality of life | x | | | | 1 | |
| Recreational mobility | | x | | | 1 | |
| Resilient societies | | | x | | 1 | |
| Sustainability | | | x | x | 2 | |
| Invasion context | | | | | | |
| Conservation values | x | | x | | 2 | |

| | | | | | | |
|---|---|---|---|---|---|---|
| Public environmental awareness (information) | x | | x | | 2 | V9,V10 |
| Purposeful introductions (Assisted colonization, etc.) | x | | | | 1 | |
| Recreation (travel) and opportunities | x | | x | | 2 | V8 |
| **iv. Technology** | | | | | | |
| General context | | | | | | |
| Artificial intelligence | | | x | | 1 | |
| Energy production | x | x | | | 2 | |
| Knowledge transfer and dissemination | | | x | | 1 | |
| Technological advances | | x | x | x | 3 | V12 |
| Invasion context | | | | | | |
| Technological advances | | | x | | 1 | |
| **v. Environment and natural resources** | | | | | | |
| General context | | | | | | |
| Agriculture | x | | | x | 2 | |
| Biodiversity | | x | x | | 2 | |
| Biofuel production | | | | x | 1 | |
| Conservation values | | | x | x | 2 | |
| Consumption of natural resources | | x | x | x | 3 | V13 |
| Ecological footprint | | x | | | 1 | |
| Environmental pollution | | x | x | | 2 | |
| Extreme/catastrophic events | x | x | | x | 3 | V14 |
| Food production | | | | x | 1 | |
| Food security | | | | x | 1 | |
| Global change/Climate change | | x | x | x | 3 | V14 |
| Landscape (urban-rural distribution; urbanization) | x | x | x | x | 4 | V17 |
| Novel ecosystems | x | | | | 1 | |
| Wilderness | | | | x | 1 | |
| Invasion context | | | | | | |
| Biodiversity | x | x | x | x | 4 | V11 |
| C sequestring crops | | | | x | 1 | |
| Crops competition | | | | x | 1 | |
| Disturbances | | | | x | 1 | |
| Global change (facilitation) | | x | | | 1 | |
| Land use (degration) | | | x | x | 2 | |
| Land use (heterogeneity, abandonment, intensificatin, etc.) | x | | x | x | 3 | V15,V16 |
| Novel ecosystems/communities | | x | | | 1 | |
| Resilience of ecosystems | | | x | | 1 | |

**Table S3.** Scoring rubric created to qualitatively assess the assumptions made under the scenarios for biological invasions for changes in: (i) 17 socio-ecological variables of interest (top rows, grouped into five different categories) and (ii) the total number of invasive alien species (IAS) (final row). Changes in variables and the number of IAS are assessed by experts using a 5-level scale ranging from -1 (high decrease) to +1 (high increase). These qualitative assessments have been associated to existing publications assessing and/or projecting changes in future trends; the use of absolute values is not intended to be exact, just to provide an overall idea of the order of change of each variable and the number of IAS across each 5-level category of change.

| VARIABLE | high increase (+1) | low increase (+0.5) | no change (0) | low decrease (-0.5) | high decrease (-1) |
|---|---|---|---|---|---|
| **POLITICS AND DEMOGRAPHICS** | | | | | |
| **V1. Democratic political culture** (based on EIU (2017)) | The pace of global democratization accelerated, supposing an important reversal in the downward tendency registered between 2006 and 2016. Regions with a recent democratic tradition consolidate their democratic developments. Most countries with authoritarian regimes shift to some sort of democratic political system, while countries with a fragile democracy become full democracies (where political freedom and civil liberties are respected, the functioning of government is satisfactory, and media and judiciary are independent). Most of the | There is a democratic transition under way in most regions worldwide. Most countries show slight improvements in some aspects of governance, political participants and media freedoms. It supposes a tipping point in the tendency registered between 2006 and 2016, although the consolidation of this democratization process is still fragile. More than half of the world's population lives in democracy of some sort, but still one-quarter live under authoritarian rules. | Democratic development remains more or less similar to 2016. North America and Western Europe present the highest democratic development, followed by Latin America. Middle East, North Africa and Sub-Saharan Africa have recorded modest improvements, while Asia has been the most successful democratizing region. One-half of the world's population lives in democracy of some sort, while one-third lives under authoritarian rules (EIU 2017). | The wave of democratization has slowed or, even, reversed. There has been a steady decline in many countries in some aspects of governance, political participation and media freedoms, continuing the tendency registered between 2006 and 2016 (EIU 2017). The biggest regressions have been in eastern Europe, North America and western Europe. Less than half of the world's population lives in democracy of some sort, while authoritarian regimes (where state political pluralism is absent and civil liberties are infringed, media is controlled by the | A process of democratic deconsolidation is under way. Most countries register important declines in some aspects of governance, political participants and media freedoms, accelerating the tendency registered between 2006 and 2016 (EIU 2017). These regressions are generalized worldwide. Most of the world's population live under authoritarian rule. |

| | | | | | |
|---|---|---|---|---|---|
| | world's population lives in democracy of some sort. | | | ruling regime and no independent judiciary) have become more common. | |
| **V2. Political globalization** | Cross-border governance plays a dominant role, enhanced by a strong partnership between governments, the private sector and civil society. International frameworks have the adequate mechanisms (political representation and collective funding via national funding co-ordination) in helping address global challenges irrespectively of national affairs. | States remains the dominant actor in national and international affairs, but there is an increasing international connectivity between a range of actors (including governments, the private sector and civil society movements). National policies are increasingly framed in global terms, reflecting the global nature of many problems and issues. | States plays the dominant role in national and international affairs, but a number of recent successes on the global governance front (the Paris COP21 and the UN's Sustainable Development Goals) show the increasing interest of states to forge international consensus on global and regional issues. The long-term maintenance of these agreements is, however, subject to major uncertainties due to the unpredictability of national affairs. | State is the dominant actor in national and international affairs, but few international frameworks have been supported by all states. The influence of international agreements on national politics is limited; national governments take decisions concerning global and regional issues based on their national priorities and expectations. | Governance at the national level, irrespectively of international affairs, is the dominant mode of governance. There are neither international agreements nor institutions. National governments take decisions concerning global and regional issues based on their national priorities and expectations, or even the interest of a few individuals. |
| **V3. Biosecurity** (prevention, eradication and control of invasive alien species (IAS), based on the prescriptions of the Aichi Biodiversity Target 9 (CBD 2010) and Honolulu Challenge on IAS (IUCN 2016)) | Most countries and islands have developed and enacted effective biosecurity policies and programs, and increased the number and scale of management actions against IAS. There is high public awareness of IAS and high public support for potential solutions. | Most countries and islands have developed and/or revised biosecurity policies and programs, but less than one-third have increased the number and scale of management actions against IAS. There are important efforts in place to raise awareness of IAS and increase public | Governments, non-governmental organizations and local communities agreed to take bold actions to counter IAS that harm biodiversity. Some countries and islands have developed and/or revised biosecurity policies and programs, but few (less than 10) | Besides the commitment of almost all of the world's countries to address IAS and some progresses developing biosecurity policies and programs in a few countries, most countries (in particular those in the developing world) have limited capacity to enforce measures to prevent | Besides the initial commitment of almost all of the world's countries to address IAS, some countries have withdrawn from international agreements. Most countries did not develop and/or revise biosecurity policies and programs. Measures to prevent future invasions and |

| | | | | | |
|---|---|---|---|---|---|
| | | support (both among relevant sectors and civil society) for potential solutions. | have committed to increase management actions against IAS. In fact, only 3% of countries are currently on track to meet these international commitments (IUCN 2016). There are some attempts to raise awareness of IAS and increase public support. | future invasions and manage existing ones. No country is currently on track to meet these international commitments. | manage existing ones have declined, as well as the resources invested in increasing public support. There is no interest in meeting these international commitments. |
| **V4. Global distribution of wealth** (considering growth, poverty and inequality, based on Hillebrand (2008)) | Very high-growth scenario, where world growth surpasses the levels registered in the last 25 years. Most countries show significant economic improvements, notably China and India, but also other regions whose recent economic history has not been favorable (such as Latin America, Africa and Middle East). This results in a substantial decrease in poverty (<353 millions of people) and a more equal Gini coefficient (<0.61, with 0 expressing perfect equality and 1 maximal inequality in wealth distribution) by 2050. | High-growth scenario, where world growth is higher than in the last 20 or 50 years. Global poverty at the $1-a-day standard falls sharply (from 965 millions of people in 2005 to 353 in 2050). Extreme poverty is eradicated in both China and India, and starts to fall in Sub-Saharan Africa. Although African economic growth rises substantially, hundreds of millions of people will remain in extreme poverty. The global Gini coefficient decreases only slightly (from 0.63 in 2005 to 0.61 in 2050), because economic growth continues to be strong in rich countries. | Moderate-growth scenario, where world growth stays the same than in the last 25 years. Global poverty at the $1-a-day standard has dropped (from 1279 millions of people in 1980 to 965 in 2005, nearing the 15% Millennium development goal). China and India accounted for most of this poverty drop, while sub-Saharan Africa registered an increase. The global Gini coefficient fell over this period from 0.65 to 0.63), but within-country distributions are less equal (mainly due to the increasing levels of inequality within China and India)(Hillebrand 2008). Ratios remain the | Low-growth scenario, where world growth does not improve (or even worsen) from the levels recorded in 1980-2005. Most countries continue the same trajectory than in the last 25 years: China and India continue their improvements, but regions that have been lagging (sub-Saharan Africa, the Middle East and Latin America) do not transition onto a high-growth path. This results in sharp increase in poverty (from 965 millions of people in 2005 to 1237 millions in 2050). The global Gini rises to 0.708 by 2050. | No-growth scenario, where world economic growth rates registered in the last 25 years become unsustainable. Most countries reduce their growth rates significantly and a few (especially those where most of the poverty is) see their growth assumptions cut drastically. This results in a dramatic increase in poverty (>1237 millions of people) and a more unequal Gini coefficient (>0.708) by 2050. |

| | | | | | |
|---|---|---|---|---|---|
| | | | same in the forthcoming decades. | | |
| **V5. Social stability** (and forced human movements, based on UNHCR (2017)) | High social and political stability, where conflicts are satisfactorily resolved in a peaceful manner (no armed conflicts). The population of forcibly displaced peoples reduces significantly, attaining values much below the ones registered in 1997 (<33.9 million). | High social and political stability, but punctuated by armed conflicts in certain parts of the world. The population of forcibly displaced people reduces substantially, attaining values similar to the ones registered in 1997 (~33.9 million) (UNHCR 2017). | Social and political stability, but unevenly distributed around the world. Conflicts are geographically and temporally localized. The population of forcibly displaced people remains similar to 2016 (~66.5 million in 2016) (UNHCR 2017). | Social and political stability combined with frequent periods of instability. Higher occurrence of armed conflicts in any part of the world. The population of forcibly displaced people grows substantially, following the trend registered over the past two decades (from 33.9 million in 1997 to 66.5 million in 2016) (UNHCR 2017). | Major social and political instability around the world, responsible of severe and continuous armed conflicts and large-scale movements of people. The population of forcibly displaced peoples increases significantly, following a more accentuated trend that the one registered over the past two decades. |
| **ECONOMY & TRADE** | | | | | |
| **V6. Trade quantity** (based on OECD/ITF (2017)) | Trade is expected to accelerate substantially, with the projected rate nearing the 5% average of the 1990s. World trade is expected to grow by a factor ≥4 between 2015 and 2050. | Trade is expected to accelerate moderately, but the projected rate will still be below the 5% average of the 1990s and subject to considerable downside risks. World trade is expected to grow by a factor of 3.5 between 2015 and 2050, which translates to a growth factor of 3.3 in tonne-kilometers for the same period (OECD/ITF 2017). | Trade is expected to accelerate slowly compared to 2016 (the fifth consecutive year of global trade growth in volume below 3%), but the projected rate will still be below the 5% average of the 1990s and subject to considerable downside risks. World trade is expected to grow by a factor of 3 between 2015 and 2050, corresponding to a growth in total tonne-kilometers (i.e. global freight transport demand) by a factor of 3.1 for the | Trade is expected to stay roughly the same or decelerate slowly, with the projected rate equal or below the 2.8% registered in 2016 [51] and subject to considerable downside risks. World trade is expected to grow by a factor ≤ 2.5 between 2015 and 2050. | Trade is expected to decelerate considerably, with the projected rate much below the 2.8% registered in 2016 [51]. World trade is expected to grow by a factor ≤ 2 between 2015 and 2050. |

| | | | | | |
|---|---|---|---|---|---|
| | | | same period (OECD/ITF 2017). | | |
| **V7. Trade distribution** | There are major shifts in the trading partners due to an increase in global trade liberalization. | There is some reorganization in trading partners, but the main patterns of global trade remain similar to today. | The main patterns of global trade remain similar. | Global trade becomes more regional. | Global trade becomes largely regional, mainly due to increased protectionism since the start of the economic crisis in 2008. |
| **V8. Passenger transport** (proxy for global connectivity, based on OECD/ITF (2017)) | Global passenger demand will more than double between 2015 and 2050, from 50000 to >120000 billion passenger-kilometers. It increases in all regions and for all transport modes, but the growth is not uniformly distributed. Developing countries (especially Asia, but also other regions) and international aviation (with 5.7% growth rate due to an increase in the air network connectivity) experience the highest growth rates, although travel demand by car will also increase the most out of all domestic non-urban modes in absolute values: an additional >45000 billion passenger-kilometres between 2015 and 2050. | Global passenger demand will double between 2015 and 2050, from 50000 to 120000 billion passenger-kilometers (prolonging the trend observed between 2010-2015). It increases in all regions and for all transport modes, but the growth is not uniformly distributed. Asia and international aviation (with 5% growth rate because the evolution of the air network remains the same than in 2010-2015) experience the highest growth rates, although travel demand by car will also increase the most out of all domestic non-urban modes in absolute values: an additional 45000 billion passenger-kilometres between 2015 and 2050 (OECD/ITF 2017). | Global passenger demand will increase between 2015 and 2050, from 50000 to <120000 billion passenger-kilometers. It increases in all regions and for all transport modes, but the growth is not uniformly distributed. Asia and international aviation (with 2.3% growth rate because the number of air connections remain the same than in 2015) experience the highest growth rates, although travel demand by car will also increase the most out of all domestic non-urban modes in absolute values: an additional <45000 billion passenger-kilometres between 2015 and 2050. | Global passenger demand will only experience a slight increase between 2015 and 2050, from 50000 to <<120000 billion passenger-kilometers. This growth is irregularly distributed among regions and for all transport modes. Asia and international aviation (with <2.3% growth rate because of changes in the number of air connections in relation to 2015) experience the highest growth rates, although travel demand by car will also experience some increase as a domestic non-urban mode of transport in absolute values: an additional <<45000 billion passenger-kilometres between 2015 and 2050. | Global passenger demand will stagnate or decrease between 2015 and 2050, oscillating around 50000 billion passenger-kilometers. This non-growth trend is irregularly distributed among regions (with Asia showing some increments), but there is a generalized deceleration in passenger demand worldwide. International aviation and travel demand by car as a domestic non-urban mode of transport remain the same than in 2015. |

| LIFESTYLE AND VALUES | | | | | |
|---|---|---|---|---|---|
| **V9. Social environmental awareness** (responsible consumption and consciousness) | There is a high environmental awareness among citizens, which empower people to participate effectively in democratic changes towards a better environment for all (e.g. reduce consumption and responsible buying). Planetary consciousness (interdependence and interconnection of all humans and the earth). | There is a high environmental awareness among citizens. People participate in changes towards a better environment for all, but their actions are not globally aligned. Consumption is reduced and responsible buying increased, but they are not an overarching priority. High consciousness of global unity. | There is environmental awareness among citizens, but their participation in changes towards a better environment is limited (e.g. lower consumption of fossil fuels, but increase in biofuels). Consumption is slightly reduced, but solutions are not always environmental friendly. Consciousness of globalism, but thinking and acting regionally. | People are aware of environmental challenges, but action is restricted to a few activities (e.g. recycle). Consumption rates remain the same. People are aware of our interconnected nature with the earth, but decisions are taken individually. | There is no environmental awareness among citizens. People act individually and unconscientiously, without caring for the environment. Selfish society. Consumption rates increase. |
| **V10. Sustainability policies** (harmonizing economic growth, social inclusion and environmental protection) | Sustainability is high in the politic agenda. The mechanisms in place are appropriate to ensure an inclusive, sustainable and resilient future for people and the planet. The Sustainable Development Goals of the 2030 Agenda for Sustainable Development (UN 2015) are reached. | Sustainability is high in the politic agenda, but the mechanisms in place are inefficient to ensure the complete achievement of the Sustainable Development Goals. There is, however, a significant advance to mitigate negative impacts. | Sustainability is in the politic agenda, but the mechanisms to ensure a desirable planet are insufficient to produce changes in past trends. There have been some significant advances in achieving the Sustainable Development Goals, but in most cases the mechanisms in place cannot reverse negative impacts. | Sustainability is rarely considered in the politic agenda. There are few mechanisms in place to ensure the accomplishment of Sustainable Development Goals and they are completely insufficient to make a significant change. | Sustainability is not a priority in the political agenda. Politicians do not care about the future of the planet and therefore continue as there were not environmental limits. |
| **V11. Status of biodiversity** | Biodiversity is highly valued and thus preserved, with environmental externalities well | Biodiversity is valued and a cost associated to ecosystem services, to account for environmental externalities. IPBES | Biodiversity is considered important, but it has only contributed to reinforce the network of protected areas. IPBES is only a | Biodiversity is known to be important, but it does not influence politic and economic decisions. IPBES does not have any | Biodiversity is not valued, neither socially nor economically. IPBES disappear and the Strategic Plan for |

| | | | | | |
|---|---|---|---|---|---|
| | integrated in the economic spheres. The influence of IPBES is high in the politic and economic agenda. The Strategic Plan for Biodiversity for the 2011-2020 was translated and implemented at the national level, and it is successively updated. | influence politics, but economic powers are of overriding importance. The Strategic Plan for Biodiversity for the 2011-2020 was translated and partially implemented at the national level. | consultation organ and the translation of the Strategic Plan for Biodiversity for the 2011-2020 was delayed at the national level. | influence and the Strategic Plan for Biodiversity for the 2011-2020 was not finally translated at the national level. | Biodiversity for the 2011-2020 was forgotten. |
| **TECHNOLOGY** | | | | | |
| **V12. Technological progress and innovation** | Higher technological progress and innovation. No limits in resources and development capacity. Technology makes significant socio-economic and environmental changes. | Higher technological progress and innovation, but geographically and socially biased. Technology makes some socio-economic and environmental changes. | Technological progress and innovation as at present, highly biased towards few countries and people. Technology influence socio-economy, but changes for environment are still limited | Technological progress and innovation similar to present, but reduced to a smaller amount of countries and people. Technology does not make significant changes in society and environment. | Lower technological progress and innovation. Limits in resources and development capacity. Technology does not have any effect on society and environment. |
| **ENVIRONMENT AND NATURAL RESOURCES** | | | | | |
| **V13. Material footprint** (proxy for pressure on natural resources, based on UNEP (2016) and Wiedmann et al. (2015)) | Pressure on natural resources accelerates and surpasses the tendencies of previous decades. Material footprint (i.e. amounts of materials that are required for production, considering both capita income and consumption + efficiency of material | Pressure on natural resources continues the increasing tendency of previous decades. Material footprint grows above 70 million metric tons (Gt) | Pressure on natural resources relent. Material footprint remains similar to 2010 (70 billion metric tons (Gt); (UNEP 2016; Wiedmann et al. 2015)). | Pressure on natural resources decelerates. Material footprint is similar to 2005 (60 billion metric tons (Gt); (UNEP 2016; Wiedmann et al. 2015)) | Pressure on natural resources exhibits a strong decrement. Material footprint is similar to 1990 (40 billion metric tons (Gt); (UNEP 2016; Wiedmann et al. 2015)). |

| | | | | | |
|---|---|---|---|---|---|
| | use) grows much above 70 billion metric tons (Gt) | | | | |
| **V14. Climate change** | High greenhouse gas concentration levels, which rise exponentially (e.g. scenario RCP 8.5; +2ºC by 2050) | Medium greenhouse gas concentration levels, which rise (e.g. scenario RCP 6 & 4.5; +1.3-1.4ºC by 2050) | Low greenhouse gas concentration levels (e.g. scenario RCP 2.6; +1ºC by 2050) | Maintenance of current greenhouse gas concentration levels, zero emissions | Reductions of current greenhouse gas concentration levels |
| **V15. Human footprint** (proxy for land degradation, based on Venter et al. (2016)) | Human footprint is widespread and intense around the world, showing a stronger increase than in 2009 (>>9%). All ecoregions show signs of human pressure and experience marked increases in their human footprints. | Human footprint is widespread and rapidly increasing around the world (>9%). Besides a few spatial divergences, most global ecoregions (>71%) show marked (>25%) increases in their human footprints. | Human footprint is widespread and rapidly increasing (9%), especially in tropical ecoregions. Wealthy and less corrupt nations show signs of improvements, yet this is overshadowed by the fact that 71% of global ecoregions show marked (>20%) increases in their human footprints. Patterns similar to the period from 1993 to 2009 (Venter et al. 2016). | Human footprint is widespread and increasing, especially in tropical ecoregions, but at a lower rate than in 2009 (<9%). Wealthy and less corrupt nations show significant improvements, yet this is overshadowed by the fact that 71% of global ecoregions show some (>10%) increases in their human footprints. | Human footprint is widespread, but decreasing thanks to the implementation of restoration measures. Most nations show significant improvements, reducing the impact of human pressure on world ecoregions (<71%). |
| **V16. Arable land** (proxy for land homogenization, based on Alexandratos and Bruinsma (2012)) | Measured from the base year 2005/2007, the net result for the world would by 2050 be an increase in the arable land area of >176 million ha, consisting of a considerable increase in the developing countries and a slight decline in the developed countries. | Measured from the base year 2005/2007, the net result for the world would by 2050 be an increase in the arable land area of 176 million ha, consisting of an increase by almost 230 million ha in the developing countries and a decline by nearly 54 million ha in the developed countries (Alexandratos and Bruinsma 2012). | Measured from the base year 2005/2007, the net result for the world would by 2050 be an increase in the arable land area of some 70 million ha, consisting of an increase by almost 110 million ha in the developing countries and a decline by nearly 40 million ha in the developed countries (as projected by | Measured from the base year 2005/2007, the net result for the world would by 2050 be an increase in the arable land area of <70 million ha, consisting of a reduced increase in the developing countries and a slight decline in the developed countries. | Measured from the base year 2005/2007, the net result for the world would by 2050 be a slight or non-increase in the arable land area, consisting of an increase in the developing countries equivalent to the decline in the developed countries. |

| | | | | | |
|---|---|---|---|---|---|
| | | | (Alexandratos and Bruinsma 2012)). | | |
| **V17. Levels of urbanization** (based on UN (2014)) | More than 80% of the world population live in urban areas, except in Africa and Asia (that present lower levels of urbanization, but a greater tendency to increase). The rate of urbanization is the same or higher than in 2015. Megacities (>10millions of inhabitants) increase worldwide. | Most people, although not the vast majority (65% approx.), live in urban areas. Urban agglomerations have increased, but the rate of urbanization has decelerated in comparison to 2015. Cities with less than 1 million inhabitants increase worldwide. | Overall, the levels of urbanization remain as in 2015 (54% in urban areas and 46% in rural areas). Medium sized-cities (around 500000 inhabitants) receive the highest attention (UN 2014). | There is a change in people's preferences from urban to rural settlements. People move to rural settlements (<5000 inhabitants), reversing the ratios from 2015 (46% in urban areas and 54% in rural areas). | There are small increments of urban agglomerations in some parts of the world, but there is a dominant tendency to live in small rural settlements (<5000 inhabitants) and disseminated houses. |
| **IAS** | **high increase (+1)** | **low increase (+0.5)** | **no change (0)** | **low decrease (-0.5)** | **high decrease (-1)** |
| **Number of established invasive alien species** | The number of established invasive alien species grows very rapidly (i.e. rates of newly established species increase, cf. Seebens et al. (2017)). | The number of established invasive alien species continues to increase rapidly (i.e. with similar rates than recently). | The number of established invasive alien species increases moderately (i.e. with slower rates than recently). | The number of established invasive alien species remains largely constant (i.e. increases only slowly and in some regions). | The number of established invasive alien species decreases (i.e. more alien species are eradicated than newly established). |

Seebens H, Blackburn TM, Dyer EE, Genovesi P, Hulme PE, Jeschke JM, Pagad S, Pysek P, Winter M, Arianoutsou M, Bacher S, Blasius B, Brundu G, Capinha C, Celesti-Grapow L, Dawson W, Dullinger S, Fuentes N, Jager H, Kartesz J, Kenis M, Kreft H, Kuhn I, Lenzner B, Liebhold A, Mosena A, Moser D, Nishino M, Pearman D, Pergl J, Rabitsch W, Rojas-Sandoval J, Roques A, Rorke S, Rossinelli S, Roy HE, Scalera R, Schindler S, Stajerova

K, Tokarska-Guzik B, van Kleunen M, Walker K, Weigelt P, Yamanaka T, Essl F (2017) No saturation in the accumulation of alien species worldwide. Nat Commun 8:14435. DOI 10.1038/ncomms14435
UN (2014) World Urbanization Prospects: The 2014 Revision (ST/ESA/SER.A/366). United Nations, New York
UN (2015) Transforming our world: the 2030 Agenda for Sustainable Development (A/RES/70/1). Unite Nations
UNEP (2016) Global material flows and resource productivity. United Nations Environment Programme, Paris
UNHCR (2017) Global trends: Forced displacement in 2016. United Nations High Commissioner for Refugees, Geneve
Venter O, Sanderson EW, Magrach A, Allan JR, Beher J, Jones KR, Possingham HP, Laurance WF, Wood P, Fekete BM, Levy MA, Watson JEM (2016) Sixteen years of change in the global terrestrial human footprint and implications for biodiversity conservation. Nat Commun 7:12558. DOI 10.1038/ncomms12558
Wiedmann TO, Schandl H, Lenzen M, Moran D, Suh S, West J, Kanemoto K (2015) The material footprint of nations. PNAS 112:6271-6276

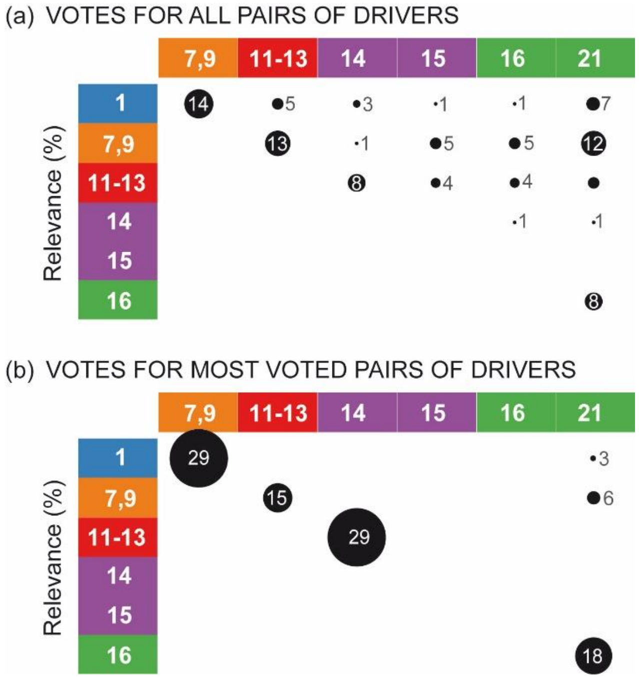


**Fig. S1.** Percentage of votes assigned by the participants of the workshop to pairs of drivers considered to develop scenarios for biological invasions. (a) shows votes assigned to all pairs of drivers in the first voting round, while (b) shows votes given in the second voting round to the six pairs of drivers presenting the highest number of votes (>5%) in (a). Columns/rows' numerical names correspond to drivers listed in Fig. 2a and the colors indicate the category of the drivers. Circles are proportional to the percentage of votes.

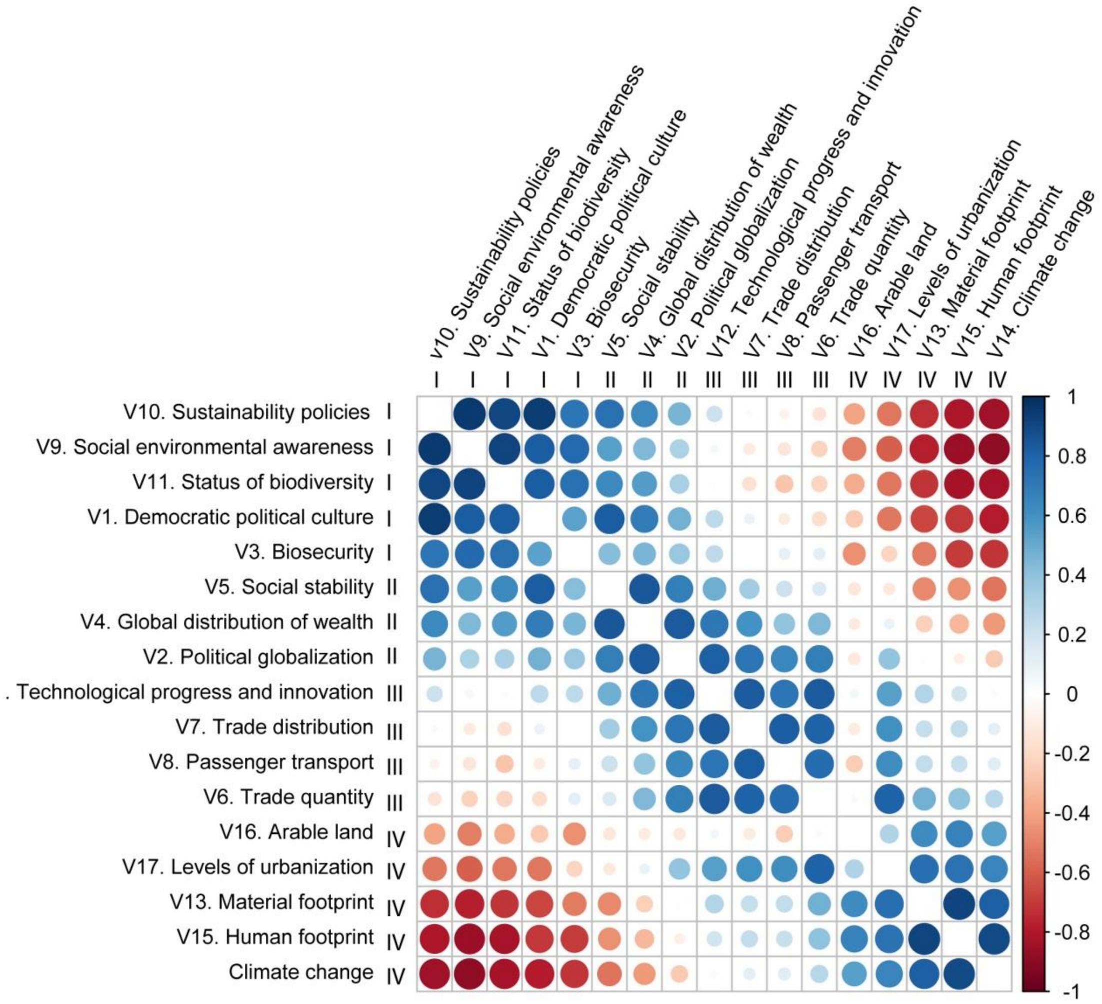


**Fig. S2.** Correlation matrix plot showing Spearman's correlation coefficient between the expert-based assessments given to socio-ecological variables of interest used to characterize scenarios for biological invasions (Fig. 4). Positive correlations are displayed in blue and negative correlations in red color; color intensity and the size of the circle are proportional to the correlation coefficients. Roman numerals correspond to the clusters of variables found in Fig. 4.
